# Chromatin-associated cohesin turns over extensively and forms new cohesive linkages during meiotic prophase

**DOI:** 10.1101/2023.08.17.553729

**Authors:** Muhammad A. Haseeb, Katherine A. Weng, Sharon E. Bickel

**Author notes:** Current address: NYU Langone Health, Mineola, NY. Corresponding author: Sharon E. Bickel.

## Abstract

In dividing cells, accurate chromosome segregation depends on sister chromatid cohesion, protein linkages that are established during DNA replication. Faithful chromosome segregation in oocytes requires that cohesion, first established in S phase, remain intact for days to decades, depending on the organism. Premature loss of meiotic cohesion in oocytes leads to the production of aneuploid gametes and contributes to the increased incidence of meiotic segregation errors as women age (maternal age effect). The prevailing model is that cohesive linkages do not turn over in mammalian oocytes. However, we have previously reported that cohesion-related defects arise in Drosophila oocytes when individual cohesin subunits or cohesin regulators are knocked down after meiotic S phase. Here we use two strategies to express a tagged cohesin subunit exclusively during mid-prophase in Drosophila oocytes and demonstrate that newly expressed cohesin is used to form *de novo* linkages after meiotic S phase. Moreover, nearly complete turnover of chromosome-associated cohesin occurs during meiotic prophase, with faster replacement on the arms than at the centromeres. Unlike S-phase cohesion establishment, the formation of new cohesive linkages during meiotic prophase does not require acetylation of conserved lysines within the Smc3 head. Our findings indicate that maintenance of cohesion between S phase and chromosome segregation in Drosophila oocytes requires an active cohesion rejuvenation program that generates new cohesive linkages during meiotic prophase.

## INTRODUCTION

During cell division, sister chromatid cohesion is required for proper alignment of chromosomes on the spindle and for accurate chromosome segregation (reviewed in: [1–6]). When sister chromatids are created during DNA replication, cohesin-mediated protein linkages are formed that keep the sisters physically associated. Within the cohesin complex, the head domains of the Smc1-Smc3 heterodimer interact with an α-kleisin subunit, forming a ring that encircles the DNA. The formation of stable cohesive linkages during S phase depends on Eco-mediated acetylation of conserved lysine residues within the head domain of the cohesin subunit, Smc3 [7–11].

In meiotic cells, sister chromatid cohesion is not only required for accurate segregation of sisters during meiosis II, but also for proper alignment and segregation of homologous chromosomes during the first meiotic division [3, 4, 6]. Within the bivalent, sister chromatid cohesion distal to the crossover keeps recombinant homologs physically associated until anaphase I. In mammalian oocytes, cohesion establishment and meiotic recombination are completed during fetal development and oocytes arrest in the dictyate stage of prophase I until ovulation. Several lines of evidence indicate that meiotic cohesion weakens as oocytes age and support the hypothesis that premature loss of cohesion in human oocytes contributes to the maternal age effect [3, 12–14]. Because turnover of cohesive linkages has not been detected in mouse oocytes [15–17], one prominent model is that human oocytes rely solely on the original cohesive linkages established before birth [3, 12–14, 18].

We have previously demonstrated that knockdown of the cohesin loader (Nipped-B), the cohesion establishment factor (Eco), or individual cohesin subunits during meiotic prophase in Drosophila oocytes leads to phenotypes consistent with premature loss of cohesion [19]. Based on these findings, we proposed that a cohesion rejuvenation program operates during meiotic prophase to establish new cohesive linkages after S phase and that this process is required to keep sister chromatids physically associated until chromosome segregation. Formation of new cohesive linkages outside of S phase is not without precedent; in mitotically dividing yeast cells, double strand breaks (DSBs) during G2 induce a genome-wide cohesion establishment program [20–22]. However, we have shown that prophase rejuvenation in Drosophila oocytes occurs even in the absence of DSBs [19], distinguishing it from the DSB-induced cohesion establishment pathway that operates in G2 yeast cells.

Using two independent approaches, we provide evidence that a tagged cohesin subunit expressed exclusively during meiotic prophase loads onto oocyte chromosomes and is used to form new cohesive linkages after meiotic S phase. Using Gal4/UAS induction to express RNAi insensitive Smc3-HA (Smc3*-HA) after meiotic S phase and Fluorescence In Situ Hybridization (FISH) to monitor cohesion, we show that expression of this transgene can rescue cohesion defects caused by knockdown of endogenous Smc3. In contrast to cohesion establishment during DNA replication, our data indicate that acetylation of conserved lysines within the Smc3 head is not required for the formation of functional cohesive linkages during meiotic prophase. In a second strategy, we created an Smc1 “tag-switch” transgene for which Gal4-induced expression of Flippase (FLP) after meiotic S phase elicits an FRT-mediated switch from mCherry-tagged to superfolder GFP-tagged Smc1 (Smc1-sGFP). Using this tool, we observe extensive turnover of chromosome-associated cohesin during meiotic prophase. Cohesin on the arms of oocyte chromosomes is completely replaced within a two-day window while turnover at centromeres is slower. Moreover, failure to load Smc1-sGFP onto chromosomes during meiotic prophase significantly increases the incidence of cohesion defects. Together our findings demonstrate that loading of cohesin and formation of new cohesive linkages after S phase are required to maintain cohesion in Drosophila oocytes during meiotic prophase.

## RESULTS

### HA-tagged Smc3* expressed after meiotic DNA replication associates with oocyte chromosomes and colocalizes with the synaptonemal complex (SC)

Here, our objective was to determine whether cohesin expressed after meiotic S phase is utilized to establish new cohesive linkages during meiotic prophase. To manipulate expression of cohesion proteins during Drosophila oogenesis, we utilized the Gal4/UAS system for both RNAi-mediated knockdown and Gal4-inducible protein expression. We designed a Gal4/UAS inducible cDNA construct to express Smc3 protein tagged with HA at its C-terminus. We engineered silent mutations in the Smc3 cDNA (denoted Smc3*) which rendered the mRNA expressed from this transgene to be insensitive to the SH00137.N hairpin used to knock down endogenous Smc3 protein (Fig 1B).

**Figure 1.**
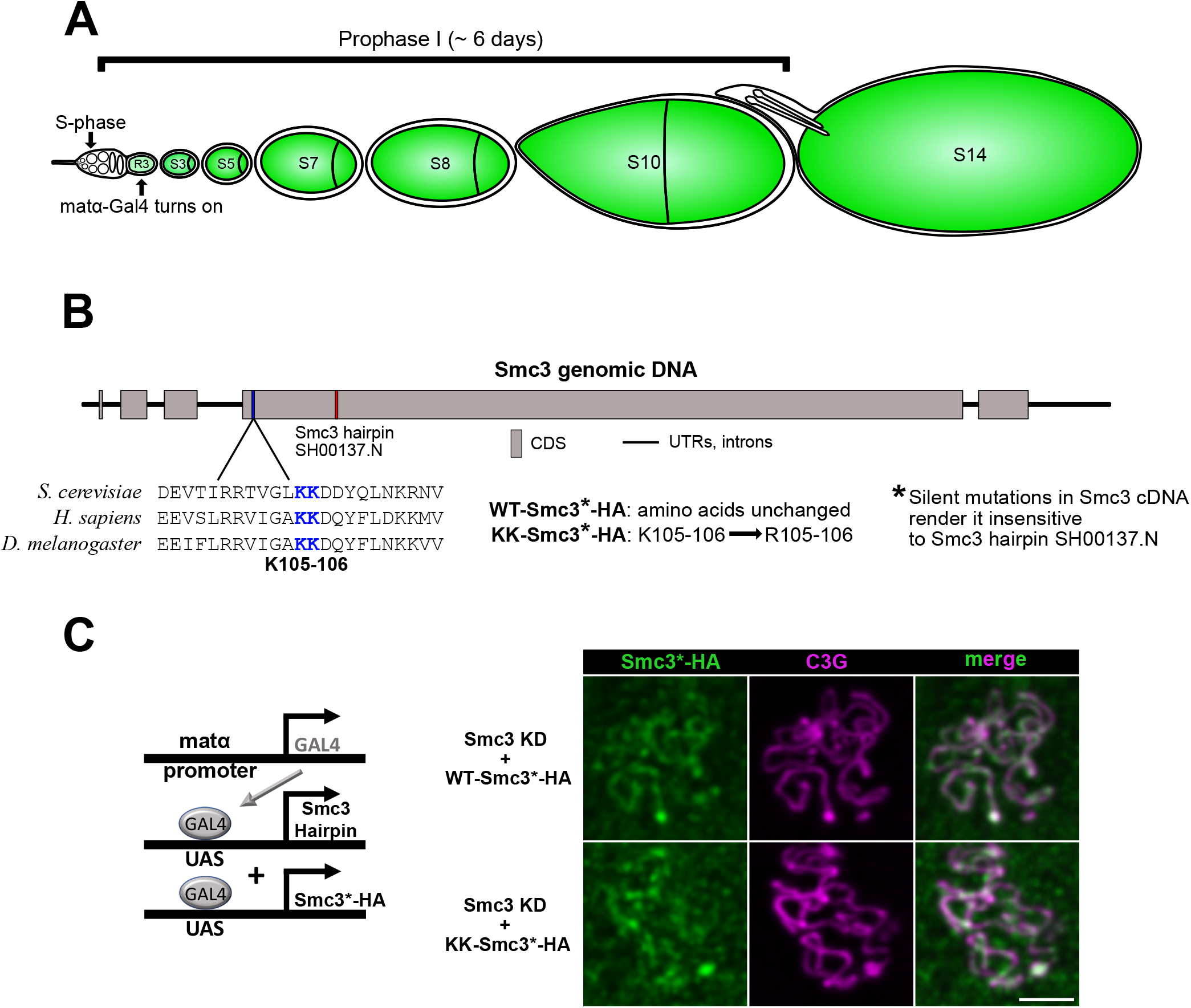
Smc3 expressed after S-phase stably associates with meiotic chromosomes independent of K105-106 acetylation. **A**. Green color depicts expression of the matα-Gal4 driver within germline cells of the Drosophila ovariole. (Note: not all stages are present in a single ovariole.) Matα-Gal4 expression begins in germarial region 3 (R3), approximately two days after meiotic S phase. Gal4 levels increase during subsequent stages. **B.** Graphic representation of the Drosophila Smc3 gene indicates the target for the short hairpin used in this study as well as the conserved lysine residues required for S-phase cohesion establishment. Silent mutations in the Smc3 cDNA (Smc3*) render transgenic Smc3*-HA constructs insensitive to the SH00137.N hairpin. Lysines 105-106 are unchanged in WT-Smc3*-HA and mutated to non-acetylatable arginines in KK-Smc3*-HA. **C. Left:** UAS-Gal4 strategy utilizes the matα-Gal4 driver to knock down endogenous Smc3 and simultaneously express HA-tagged Smc3* protein (hairpin insensitive) during meiotic prophase. **Right:** matα-Gal4 induced knockdown of endogenous Smc3 and expression of WT-Smc3*-HA or KK-Smc3*-HA during meiotic prophase results in HA signal that colocalizes with the synaptonemal complex protein, C(3)G. Stage 2 oocytes are shown. Images are maximum intensity projections of deconvolved confocal Z series. Scale bar, 2μm.

The matα-Gal4-VP16 driver (hereafter named matα-Gal4) is germline specific and turns on approximately two days after exit from meiotic S phase [19], specifically during mid-prophase (see Fig 1A). Use of this driver allows us to modify protein levels after the original cohesive linkages have already been established during meiotic S phase. Using this strategy, we have previously demonstrated that matα-Gal4 induced knockdown of cohesion proteins during meiotic prophase in Drosophila oocytes causes phenotypes indicative of premature loss of cohesion [19]. Here, we utilize “strong” and “weak” matα-Gal4 driver chromosomes, which we have previously described [23]. These driver chromosomes also contain an amorphic allele of the *matrimony* gene (*mtrm^KG0805^)* [24].

Using the strong matα-Gal4 driver to knock down endogenous Smc3 and simultaneously express wild-type Smc3 tagged with HA (WT-Smc3*-HA), we asked whether HA-tagged Smc3* could associate with the meiotic chromosomes during prophase (Fig 1C). Following HA immunolocalization, we observed long continuous threads of chromatin-associated WT-Smc3*-HA signal in the oocyte starting in Region 3 (R3) of the germarium, the stage at which the matα-Gal4 driver is first expressed [19]. Cohesin-enriched chromosome axes form a scaffold for assembly of the synaptonemal complex (SC), a multi-protein structure that holds homologous chromosomes in close proximity during meiotic recombination [25–28]. WT-Smc3*-HA localization overlaps extensively with that of the SC protein C(3)G (Fig 1C) and mimics what we have previously described for endogenous cohesin subunits Smc1 and Smc3 [29]. These data indicate that a cohesin subunit synthesized after meiotic S phase associates with the meiotic chromosomes during prophase.

### SC defects caused by prophase knockdown of endogenous Smc3 protein are suppressed by simultaneous expression of HA-tagged-Smc3*

To assess whether WT-Smc3*-HA protein that loads onto chromosomes after meiotic S phase becomes part of functional cohesin complexes, we asked whether SC defects caused by prophase knockdown of endogenous Smc3 could be suppressed by simultaneous expression of HA-tagged Smc3*, which is RNAi insensitive. We have previously shown that matα-Gal4 driven knockdown of individual cohesion proteins causes premature disassembly of the SC as early as stage 2 [19]. Therefore, SC defects serve as a proxy for premature loss of meiotic cohesion. In wild-type Drosophila oocytes, the SC fully assembles in germarial region 2A and usually remains intact until stages 5-6, although the timing of disassembly can be affected by genetic background [30, 31].

To quantify SC defects, we visualized C(3)G immunostaining and assigned each oocyte to one of four categories (Fig 2A). Long continuous threads of C(3)G signal are indicative of normal, full-length SC. Broken threads, short threads, and spots correspond to SC defects with increasing severity. For control oocytes (Fig 2B), we utilized females that lacked a Gal4 driver but contained the UAS-Smc3 hairpin transgene as well as an “Empty” UASp vector insertion that does not code for protein. Inclusion of this “UAS-Empty” transgene allowed us to match the number of UAS binding sites in control and experimental genotypes and avoid confounding results due to Gal4 titration effects. We observed full-length SC in all control oocytes until stage 6, at which time a few oocytes had begun normal disassembly of the SC (Fig 2B).

**Figure 2.**
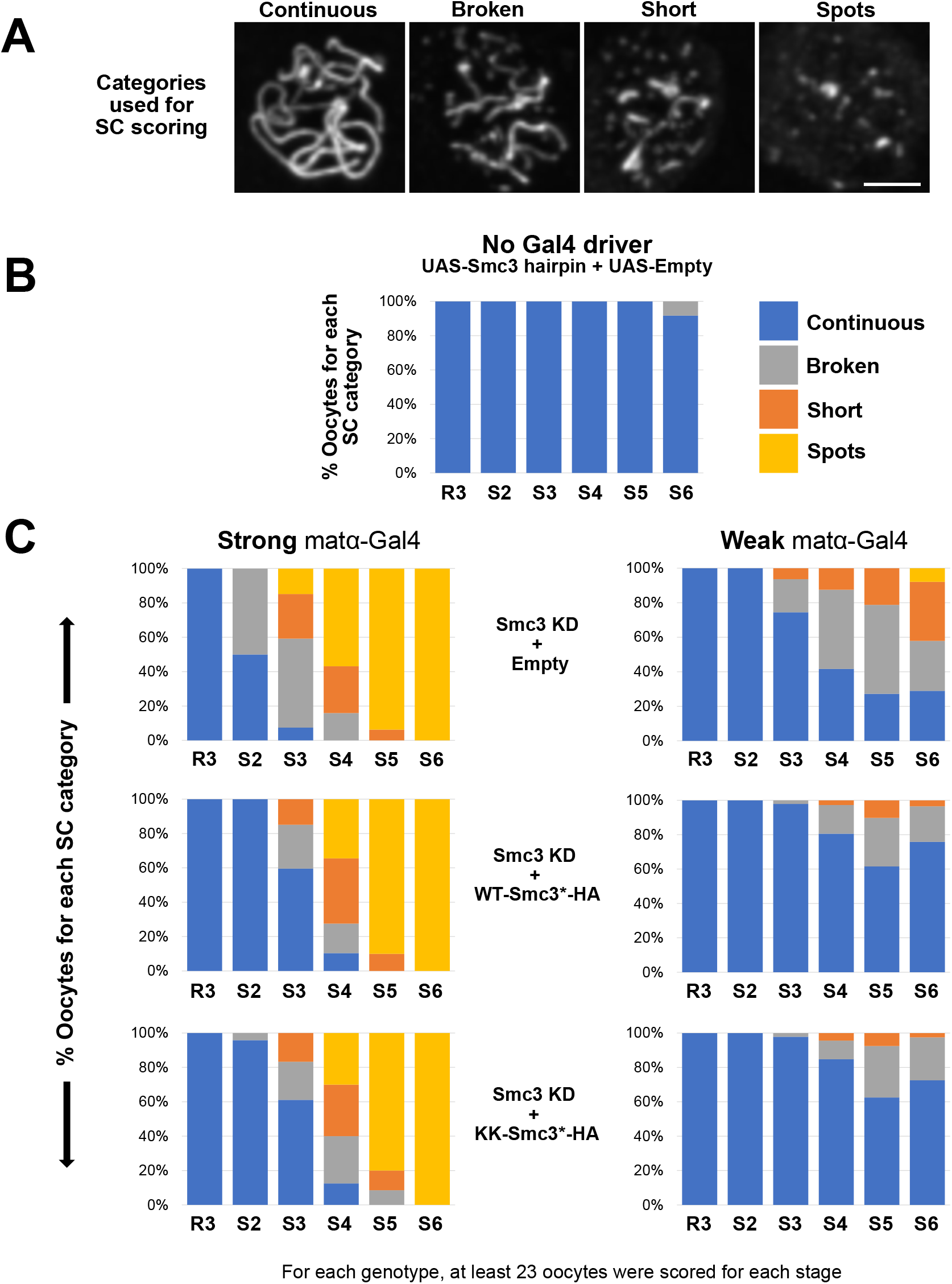
Prophase expression of HA-tagged-Smc3* delays premature SC disassembly caused by knockdown of endogenous Smc3 protein. **A**. The integrity of the synaptonemal complex (SC) was assessed using C(3)G immunostaining and oocytes were assigned to one of the four categories shown. “Continuous” corresponds to normal SC with long continuous threads. Premature disassembly was categorized as “Broken” (long fragments), “Short” (short fragments) or “Spots” (the most severe phenotype). Images are maximum intensity projections of deconvolved confocal Z series. Scale bar, 2μm. **B.** SC defects were quantified in oocytes from region 3 (R3) to stage 6 (S6). In the control genotype, SC disassembly was not detected until stage 6. Because these flies lack a Gal4 driver, the UAS-Smc3 hairpin and UAS-Empty transgene are not expressed. Persistence of full-length SC until stage 6 is normal in wild-type flies. **C. Left:** When the strong matα-Gal4 driver is utilized to knock down endogenous Smc3, SC defects are visible in stage 2 with severe defects evident by stage 3. However, premature disassembly of the SC is delayed when WT-Smc3*-HA or KK-Smc3*-HA protein expression is induced during prophase in Smc3 KD oocytes. **Right:** SC defects are milder when the weak matα-Gal4 driver is used to knock down endogenous Smc3. These defects are delayed and less severe when Smc3*-HA is simultaneously expressed with the Smc3 hairpin during meiotic prophase.

When the strong matα-Gal4 driver is combined with the UAS-Smc3 hairpin and UAS-Empty vector (Smc3 KD + Empty), SC defects are visible in stage 2 and increase in severity in subsequent stages (Fig 2C, left). These results are very similar to what we previously observed for matα-Gal4 driven expression of the same Smc3 hairpin in the absence of UAS-Empty [19]. In contrast, when expression of WT-Smc3*-HA and knockdown of endogenous Smc3 are simultaneously driven by strong matα-Gal4 (Smc3 KD + WT-Smc3*-HA), SC defects do not arise until stage 3 and are less severe in stage 4 than for Smc3 KD oocytes lacking HA-tagged Smc3* (Fig 2C left). This result supports the hypothesis that cohesin complexes that load onto meiotic chromosomes during prophase are functional. However, expression of WT-Smc3*-HA only marginally reduces the frequency and severity of SC defects in stage 5 oocytes (compared to Smc3 KD + Empty), and SC defects in stage 6 are equivalent in the two genotypes (Fig 2C left). We reasoned that SC defects in later stages might arise if overexpression of WT-Smc3*-HA caused by the strong matα-Gal4 driver perturbs the normal stoichiometry of cohesin subunits and disrupts proper cellular localization of cohesin as described for cultured mammalian cells [32]. Consistent with this idea, WT-Smc3*-HA induced by strong matα-Gal4 is primarily cytosolic during stages 5 and 6, with little to no nuclear signal (Fig S1).

To express lower levels of RNAi insensitive HA-tagged Smc3* specifically during prophase, we utilized a previously described weak matα-Gal4-VP16 driver chromosome [23]. As expected, expression of the Smc3 hairpin with the weak driver resulted in SC defects that were not as severe and began later than those observed with the strong driver (compare left and right top graphs in Fig 2C). When we induced expression of both WT-Smc3*-HA and the Smc3 hairpin with the weak matα-Gal4 driver, premature disassembly of the SC started later than in Smc3 KD + Empty oocytes and the defects were mild (compare left and right middle graphs in Fig 2C), again supporting the hypothesis that cohesin loaded onto meiotic chromosomes during prophase is functional.

Neither the strong nor weak matα-Gal4 driver chromosome results in Smc3*-HA expression that completely rescues the SC defects caused by knockdown of endogenous Smc3. However, partial suppression with each of these drivers argues that the cohesin we observe loading onto meiotic chromosomes during prophase is functional (Fig 2C). The level of prophase-expressed HA-tagged Smc3* relative to endogenous cohesin subunits is likely critical for its functionality [32]. Although we have been unable to detect Smc3*-HA signal above background at any stage when using the weak driver, SC defects in stages 5 and 6 are better suppressed when the weak driver is used. These data support the hypothesis that mislocalization of WT-Smc3*-HA caused by strong matα-Gal4 driver induced overexpression is responsible for the severe SC defects we observe in stages 5-6 in that genotype.

### HA-tagged Smc3* expressed during meiotic prophase is used to establish new cohesive linkages

To directly assay the state of sister chromatid cohesion in Smc3 KD oocytes expressing low levels of RNAi insensitive WT-Smc3*-HA with the weak matα-Gal4 driver, we utilized Fluorescence In Situ Hybridization (FISH). Using the 359-bp satellite repeat probe that binds to *X* chromosome pericentric heterochromatin and an Oligopaint arm probe that hybridizes to a distal 100kb region on the *X* chromosome (Fig 3A), we quantified cohesion defects at these two locations in mature Drosophila oocytes (stages 13 and 14). Sister chromatid cohesion distal to the crossover keeps recombinant homologs physically associated (Fig 3A), and premature loss of arm cohesion results in segregation errors during the first meiotic division. Using a different arm probe, we have previously reported an increase in arm cohesion defects, but not pericentric cohesion defects, when the original matα-Gal4 driver chromosome (*mtrm^+^*) is used to induce expression of the SH00137.N Smc3 hairpin [33].

**Figure 3.**
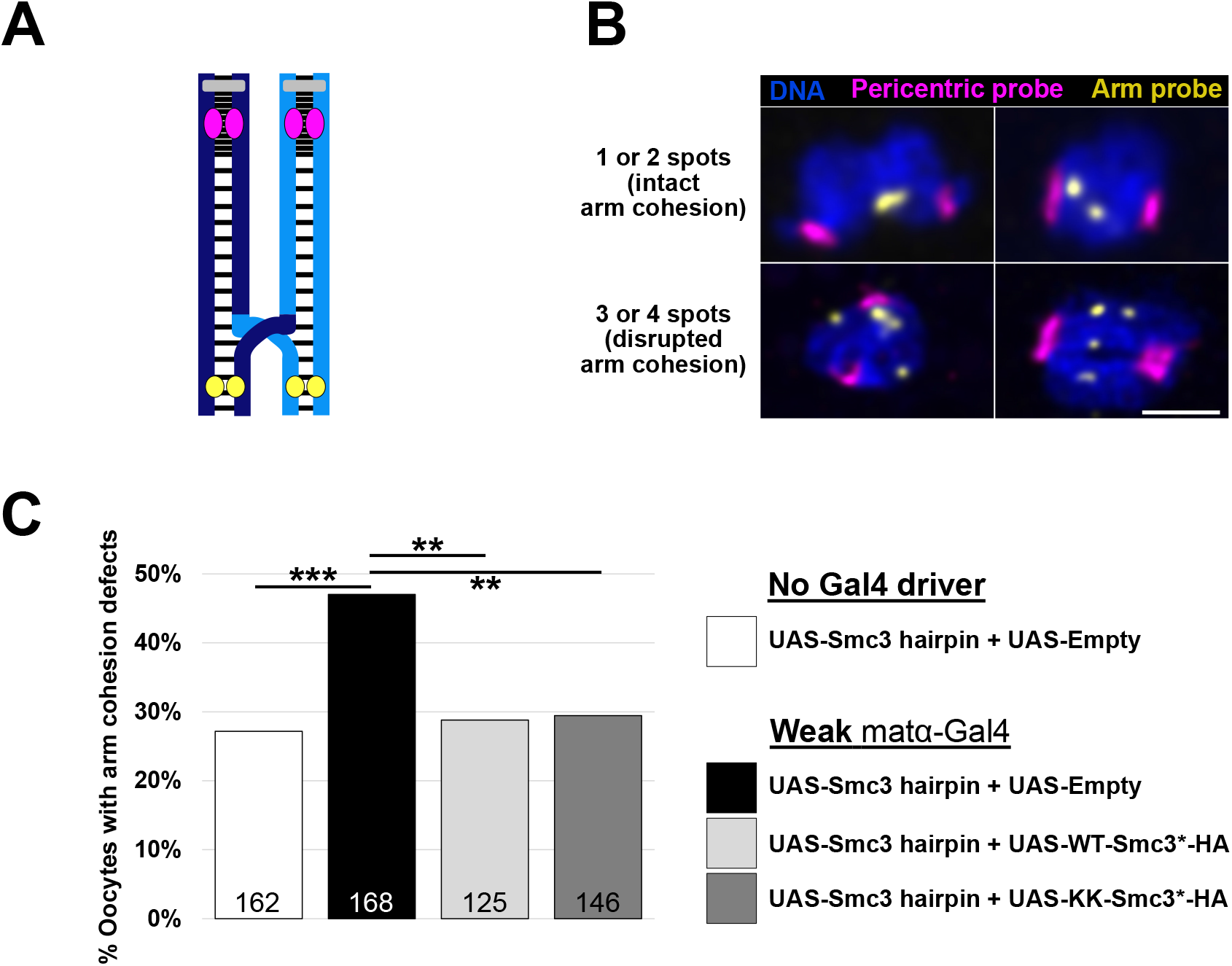
New cohesive linkages are established during meiotic prophase and their formation does not require acetylation of Smc3 at conserved lysines K105-106. **A**. Schematic depicts a recombinant X chromosome bivalent in Drosophila oocytes. Homologous chromosomes (each composed of two sister chromatids) are shown in dark blue and light blue with centromeres in grey. Cohesion between sister chromatids, depicted as black lines, keeps the recombinant homologs associated. Magenta and yellow represent the regions to which pericentric heterochromatin and Oligopaint arm probes hybridize, respectively. **B.** Representative images are shown for intact arm cohesion (1 or 2 arm probe spots) and premature loss of arm cohesion (3 or 4 arm probe spots, quantified as arm cohesion defects). Images are maximum intensity projections of deconvolved confocal Z series. Scale bar, 2μm. **C.** Arm cohesion defects in mature oocytes are quantified for each genotype shown. Number of oocytes scored for each genotype is shown within each bar. A two-tailed Fisher’s exact test was used to calculate P values. *** corresponds to P < 0.001 and ** corresponds to P < 0.002.

We first quantified cohesion defects in oocytes that contained both the UAS-Smc3 hairpin and the UAS-Empty transgene, comparing genotypes with or without the weak matα-Gal4 driver. We considered arm cohesion intact if the arm probe resulted in either one or two spots. Those oocytes with three or four arm probe spots were counted as arm cohesion defective (Fig 3B). Expression of the Smc3 hairpin induced by the weak matα-Gal4 driver caused a significant increase in the percentage of oocytes exhibiting arm cohesion defects compared to those lacking a Gal4 driver (Fig 3C, P < 0.001). Pericentric cohesion defects were negligible in Smc3 KD oocytes, consistent with our previous studies [33]. Our results demonstrate that even the weak matα-Gal4 driver results in Smc3 knockdown during meiotic prophase that is sufficient to cause premature loss of arm cohesion in prometaphase and metaphase I arrested Drosophila oocytes.

We then asked whether expression of RNAi insensitive WT-Smc3*-HA after meiotic S phase can suppress arm cohesion defects caused by knockdown of endogenous Smc3. When the weak matα-Gal4 driver was used to express both the Smc3 hairpin and the WT-Smc3*-HA transgene during meiotic prophase, the number of oocytes with premature loss of arm cohesion was significantly lower than for Smc3 KD + UAS-Empty oocytes (Fig 3C, P < 0.002). Moreover, the incidence of defects in Smc3 KD oocytes expressing WT-Smc3*-HA was similar (P = 0.79) to that in control oocytes that lack a Gal4 driver (no knockdown or protein expression), consistent with complete rescue. These data demonstrate that new cohesive linkages are established after meiotic S phase in Drosophila oocytes.

### Formation of new cohesive linkages during meiotic prophase does not require acetylation of Smc3 at conserved lysines 105-106

During DNA replication, the formation of stable cohesive linkages depends on Eco-mediated acetylation of conserved lysines within the Smc3 head domain [7–11]. These lysines correspond to K105-106 in Drosophila Smc3 (Fig 1B). We have previously demonstrated that matα-Gal4 induced knockdown of the acetyltransferase Eco in Drosophila oocytes causes premature disassembly of the SC and chromosome segregation errors consistent with premature loss of arm cohesion [19]. These results support the hypothesis that acetylation of one or more targets by Eco is required for cohesion rejuvenation in Drosophila oocytes during meiotic prophase.

To determine whether acetylation of lysines 105-106 in Drosophila Smc3 is required for the formation of functional cohesive linkages during meiotic prophase, we utilized a second RNAi insensitive HA-tagged Smc3* cDNA construct in which these lysines are mutated to non-acetylatable arginine residues (KK-Smc3*-HA, Fig 1B). When expression of KK-Smc3*-HA was induced by the strong matα-Gal4 driver, we observed threads of HA signal colocalizing with the SC protein C(3)G (Fig 1C), indicating that acetylation of K105-106 is not required for association of Smc3 with oocyte chromosomes during meiotic prophase. Furthermore, KK-Smc3*-HA expressed during meiotic prophase was able to function as well as WT-Smc3*-HA in its ability to suppress SC defects caused by knockdown of endogenous Smc3 (Fig 2C). This was true whether we utilized the strong or weak matα-Gal4 driver chromosome to induce simultaneous knockdown and protein expression (compare middle and lower panels in Fig 2C). Finally, like WT-Smc3*-HA, expression of KK-Smc3*-HA during meiotic prophase rescued arm cohesion defects caused by Smc3 knockdown (Fig 3C, P < 0.002). These data indicate that formation of new cohesive linkages in Drosophila oocytes during meiotic prophase does not depend on Eco-dependent acetylation of Smc3 at K105-106.

### Chromatin-associated cohesin turns over extensively during meiotic prophase

One drawback of matα-Gal4 induction of UAS-Smc3*-HA during meiotic prophase is that we have been unable to express HA-tagged Smc3* at a level comparable to endogenous Smc3. In addition, we wanted to rule out the possibility that a cohesin subunit expressed during prophase associates with chromatin and contributes to formation of new cohesive linkages only if we knock down the endogenous subunit.

To address the above concerns, we designed a construct that allowed us to visualize tagged-Smc1 protein expressed at physiological levels specifically during meiotic prophase (Smc1 tag-switch transgene, Fig 4A). This transgene, integrated at the attP40 site on chromosome 2, is composed of Smc1 genomic DNA, including upstream regulatory sequences that control its expression. In addition, the construct encodes a switchable fluorescent protein tag fused to the C-terminus of the encoded Smc1 protein (Fig 4A). In the absence of Flippase (FLP) expression, Smc1 encoded by this transgene is tagged with mCherry. However, because the mCherry coding sequence (including a stop codon) is flanked by FRT sites, induction of FLP expression causes excision of mCherry DNA and results in expression of Smc1 protein fused to superfolder-GFP (sGFP) [34].

**Figure 4.**
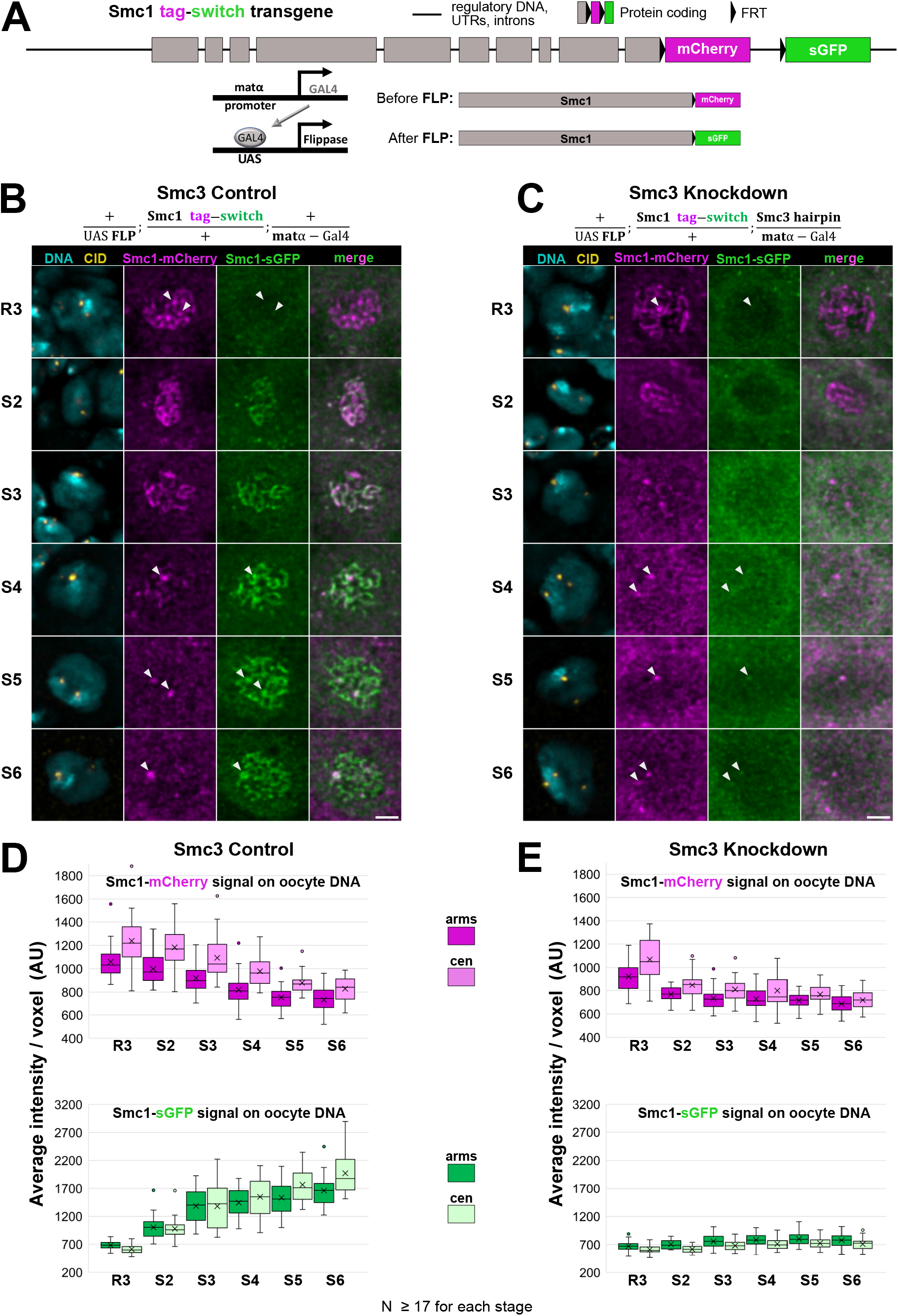
Chromosome-associated cohesin turns over extensively during meiotic prophase. **A**. Details of the Smc1 tag-switch transgene and expression strategy are illustrated. The construct, inserted into the attP40 site on chromosome 2, utilizes Smc1 genomic DNA with expression controlled by *smc1* regulatory sequences. The mCherry fluorescent tag is flanked by FRT sites. Flippase (FLP) expression induced by the original matα-Gal4 driver chromosome causes excision of the mCherry tag and expression of Smc1 tagged with superfolder-GFP (sGFP). **B-C.** Representative Smc1 tag-switch images are presented for region 3 (R3) through stage 6 (S6) oocytes in which Smc3 is unperturbed (Smc3 control, **B**) or knocked down during prophase (Smc3 Knockdown, **C**). Oocyte DNA (cyan), CID (yellow), Smc1-mCherry (magenta) and Smc1-sGFP (green) are shown. Arrowheads mark the location of centromeres (based on CID signal). Images are maximum intensity projections of deconvolved confocal Z series. For each fluorescent tag, images were acquired and processed identically for both genotypes. Scale bars, 2μm. **D-E.** Box and whisker plots quantify signal intensity for Smc1-mCherry (magenta) and Smc1-sGFP (green) on chromosome arms (dark) and centromeres (light) at the indicated oocyte stages in Smc3 control oocytes (**D**) and Smc3 knockdown oocytes (**E**). The average intensity is indicated by an “X”, the median and quartiles depicted with horizontal lines. AU = arbitrary units.

Using Smc1 immunoblotting, we verified that the level of tagged Smc1 protein produced from the tag-switch transgene is similar to that encoded by a single copy of the endogenous *smc1* gene (Fig S2A). In addition, we confirmed that the transgene rescues the lethality of *smc111* homozygotes (Fig S2B), indicating that the tagged Smc1 protein is functional.

We utilized the original matα-Gal4 driver chromosome (*mtrm*^+^) to induce expression of a UAS-FLP transgene in the germline of females also containing the Smc1 tag-switch insertion (Fig 4B). In this genotype, the onset of FLP expression in germarial region 3 (see Fig 1A) should elicit a switch from Smc1-mCherry to Smc1-sGFP expression, allowing us to visualize sGFP-tagged Smc1 produced exclusively during meiotic prophase.

Following nanobody staining, we monitored the localization and signal intensity of chromosome-associated Smc1-mCherry and Smc1-sGFP in oocytes from germarial region 3 (R3) to stage 6 (S6). The expected pattern of long continuous Smc1-mCherry threads on oocyte DNA was evident in R3 (Fig 4B) as well as earlier stages within the germarium (Fig S3). In contrast, we did not observe distinct Smc1-sGFP signal on oocyte chromosomes until stage 2 (Fig 4B & Fig S3), and its intensity was relatively weak (Fig 4B & 4D). During subsequent stages, the signal for sGFP-tagged Smc1 became more prominent on both the chromosome arms and the centromeres (Fig 4B & 4D). In stages 2 and 3, we observed extensive colocalization of the mCherry and sGFP signals on oocyte DNA. However, as the signal intensity for chromosomal sGFP-tagged Smc1 increased, we observed a concomitant decrease in Smc1-mCherry associated with the meiotic chromosomes (Fig 4B & 4D). Interestingly, mCherry-tagged Smc1 signal persisted longer on the centromeres than on the chromosome arms (Fig 4D). In stages 5 and 6, we observed both Smc1-sGFP and Smc1-mCherry at oocyte centromeres, but distinct threadlike signal on the arms was only apparent for sGFP-tagged Smc1 (Fig 4B & 4D). These data indicate that chromosome-associated cohesin turns over extensively during meiotic prophase. Importantly, association of prophase-specific Smc1-sGFP with meiotic chromosomes occurs even when the level of endogenous Smc1 is not perturbed by knockdown.

### Failure to load cohesin onto meiotic chromosomes during meiotic prophase leads to premature loss of arm cohesion

To further characterize Smc1 tag-switch dynamics during meiotic prophase, we examined Smc1-mCherry and Smc1-sGFP localization in oocytes in which expression of the UAS-Smc3 hairpin was induced by the original matα-Gal4 driver chromosome (see genotype, top of Fig 4C). When Smc3 was knocked down, loading of newly synthesized sGFP-tagged-Smc1 onto oocyte chromosomes was almost completely abolished (Fig 4C & 4E), although at later stages we detected faint Smc1-sGFP signal at some centromeres (Fig 4C, arrowheads in S4-S6). As we observed for Smc3 control oocytes, chromosome-associated Smc1-mCherry decreased during prophase progression in Smc3 KD oocytes, but the decline was more precipitous than in control (compare Fig 4D & 4E). However, mCherry-tagged Smc1 was still visible at oocyte centromeres at later stages in Smc3 KD oocytes (Fig 4C, arrowheads in S4-S6). In contrast to Smc3 control oocytes (Fig 4B), Smc1-sGFP and Smc1-mCherry signals were often most predominant in the cytoplasm, not the nuclei, of Smc3 KD oocytes (Fig 4C). Based on these data, we conclude that loading of newly synthesized sGFP-tagged-Smc1 onto chromosomes during meiotic prophase depends on the other cohesin subunits and that the Smc1-sGFP dynamics that we observe reflect the behavior of the intact cohesin ring.

We also examined Smc1 tag-switch localization in stage 10 oocytes for both Smc3 control and knockdown genotypes. This late prophase stage occurs approximately three days after the onset of matα-Gal4 expression [19, 35]. We have been unable to detect endogenous or tagged-cohesin subunits on the condensed chromosomes of metaphase I arrested oocytes (stage 14), most likely due to technical limitations. However, in well-fed females undergoing normal egg-laying, it takes only a short time (∼three hours) for a stage 10 oocyte to mature to stage 14 and arrest at metaphase I [35].

In the presence of normal levels of Smc3 (Smc3 control), sGFP-tagged Smc1 was enriched in the large nucleus of the stage 10 oocyte (Fig S4A-B) and associated with the meiotic chromosomes (Fig 5A). In contrast, when Smc3 was knocked down, Smc1-sGFP was largely excluded from the nuclei of stage 10 oocytes (Fig S4C-D) and failed to localize to the meiotic chromosomes (Fig 5A-B). Notably, even in late prophase stage 10 oocytes, mCherry-tagged Smc1 was still visible at the centromeres in both genotypes (Fig 5A, arrowheads).

**Figure 5.**
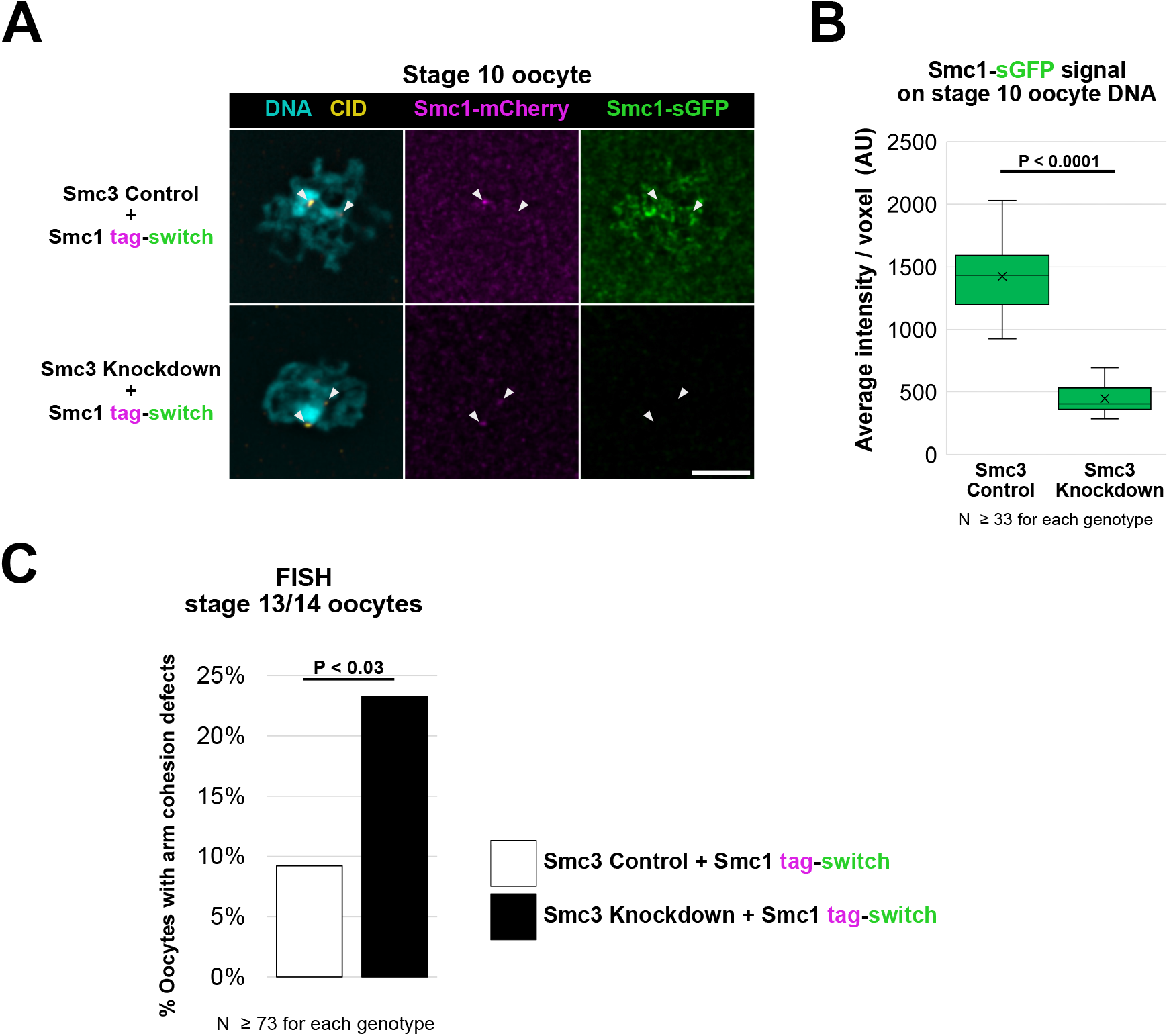
Failure to load newly synthesized cohesin on oocyte chromosomes during meiotic prophase leads to premature loss of arm cohesion. **A**. Chromatin localization for Smc1-mCherry and Smc1-sGFP during stage 10 is shown for Smc3 control and Smc3 knockdown oocytes. Images are maximum intensity projections of deconvolved confocal Z series. For each fluorescent tag, images were acquired and processed identically for both genotypes. Scale bar, 2μm. **B.** Chromatin-associated Smc1-sGFP signal intensity during stage 10 is quantified for Smc3 control and Smc3 knockdown oocytes. The average intensity is indicated by an “X”, the median and quartiles are depicted with horizontal lines. AU = arbitrary units. **C.** FISH was used to quantify arm cohesion defects in Smc3 control and Smc3 knockdown oocytes expressing the Smc1 tag-switch transgene. P value was calculated using a two-tailed Fisher’s exact test.

To ask whether chromosome-associated Smc1-sGFP participates in the formation of new cohesive linkages during meiotic prophase, we performed FISH to quantify cohesion defects in Smc3 control and knockdown oocytes expressing the Smc1 tag-switch transgene (Fig 5C). In Smc3 knockdown oocytes, in which Smc1-sGFP fails to load onto the oocyte DNA during meiotic prophase, cohesion defects were significantly higher than in Smc3 control oocytes (Fig 5C, P < 0.03). These data indicate that normal turnover and loading of cohesin complexes during meiotic prophase leads to the formation of new cohesive linkages that are required to maintain arm cohesion in Drosophila oocytes.

## DISCUSSION

Using two different approaches, we have observed that cohesin synthesized after meiotic S phase can load onto Drosophila oocyte chromosomes and facilitate the formation of new cohesive linkages during prophase. When we use the prophase-specific matα-Gal4 driver to knock down endogenous Smc3, simultaneous expression of a UAS-Smc3*-HA transgene can partially suppress knockdown-induced SC defects (Fig 2) and fully rescue arm cohesion defects (Fig 3). However, one challenge with this approach is achieving physiological levels of Smc3*-HA expression. Therefore, we used an alternative strategy in which genomic sequences within an Smc1 transgene resulted in expression of tagged Smc1 protein at levels that were comparable to the endogenous protein. Matα-Gal4 induced expression of FLP during meiotic prophase causes a switch from Smc1-mCherry to Smc1-sGFP expression from this transgene, allowing us to follow the behavior of a cohesin subunit synthesized exclusively during prophase. Using this Smc1 tag-switch construct, we observed nearly complete turnover of cohesin on oocyte chromosomes within a two-day timeframe (R3 to S6, Fig 4B & 4D). Given that meiotic prophase lasts approximately six days in Drosophila oocytes [35], our data suggest that chromatin-bound cohesin complexes could be replaced three times during the course of meiosis. Importantly, when Smc3 is knocked down, chromosome association of Smc1-sGFP during meiotic prophase is nearly abolished (Fig 4C & 4E). Moreover, failure of sGFP-tagged Smc1 to associate with the meiotic chromosomes during meiotic prophase results in premature loss of arm cohesion (Fig 5). These data provide compelling evidence that turnover of cohesin on meiotic chromosomes is normal in Drosophila oocytes and leads to the formation new cohesive linkages after DNA replication.

Our previous work indicated that cohesion rejuvenation during meiotic prophase requires the acetyltransferase Eco [19]. During DNA replication, cohesion establishment depends on Eco-mediated acetylation of two lysines within the Smc3 head domain [7–11]. In contrast, we show here that formation of new linkages during meiotic prophase does not require acetylation of these conserved lysines (K105-6) within Drosophila Smc3; when endogenous Smc3 is knocked down, expression of KK-Smc3*-HA rescues arm cohesion defects as well as WT-Smc3*-HA (Fig 3, P = 0.99). Our data suggest that formation of stable cohesive linkages during meiotic prophase depends on Eco-dependent acetylation of other lysines within Smc3 or a completely different protein target. In mitotically dividing yeast cells, Eco acetylation of the mitotic α-kleisin (not Smc3) is required for genome-wide cohesion establishment during G2 in response to DNA damage [36, 37]. We have previously shown that when meiotic double strand breaks (DSBs) are prevented (Spo11 null oocytes), Eco is still required during meiotic prophase to prevent cohesion-related defects, indicating that the cohesion rejuvenation pathway in Drosophila oocytes does not depend on induction of DSBs [19]. Therefore, formation of new cohesive linkages during meiotic prophase in Drosophila oocytes is mechanistically distinct from cohesion establishment during S phase and DNA-damage induced cohesion establishment during G2 in yeast cells. Future experiments will be required to identify the Eco target required for cohesion rejuvenation during meiotic prophase.

The data we present here indicate that association of cohesin with chromosomes is highly dynamic during meiotic prophase in Drosophila oocytes. Interestingly, our observation that Smc1-mCherry turnover at centromeres is slower than that on arms suggests that cohesive linkages within pericentric heterochromatin are more stable than those on chromosome arms. This may be why we do not detect pericentric cohesion defects with the 359 bp satellite DNA probe when we knock down Smc3 (this paper, and [33]). Alternatively, the large chromosomal target for this probe (∼11Mb) may impede our ability to detect cohesion defects in pericentric heterochromatin.

Our experiments expressing UAS-Smc3*-HA utilized matα-Gal4 driver chromosomes (strong and weak) that also carried an amorphic allele of the *matrimony* gene (*mtrm^KG08051^,* abbreviated *mtrm^KG^*). In Drosophila oocytes, accurate segregation of achiasmate homologs relies on homology-dependent interactions within their pericentric heterochromatin [38, 39], and heterozygosity for *mtrm^KG^* causes missegregation of achiasmate homologs [24]. Because the achiasmate segregation system also promotes accurate segregation of recombinant homologs that have lost chiasmata due to premature loss of arm cohesion, we often utilize a *mtrm ^KG^/+* background for chromosome segregation assays to disable this mechanism [23, 33, 40]. Our previous creation of recombinant chromosomes that contain both the matα-Gal4-VP16 transgene and the *mtrm ^KG^* allele resulted in driver chromosomes of different strengths [23], which proved useful for this study.

In our FISH analyses, the baseline for arm cohesion defects in control oocytes was considerably higher in *mtrm ^KG^/+* oocytes (Fig 3) than in oocytes with two wild-type copies of *mtrm* (Fig 5). Mtrm protein binds and inhibits the activity of Drosophila Polo-like kinase (PLK) in a dose dependent manner [41]. Furthermore, in both mitotic and meiotic cells, PLK phosphorylation of cohesin has been shown to cause its dissociation from chromosome arms during prophase in a cleavage-independent manner [42–44]. *mtrm* null oocytes exhibit a high incidence of completely separated chromatids [45], consistent with elevated PLK activity in late prophase inducing premature dissociation of cohesin from the meiotic chromosomes. However, in *mtrm* heterozygotes, cohesion defects were limited to the distally translocated heterochromatin of *FM7* in the *FM7/X* achiasmate homolog pair and negligible for *FM7/FM7* or *X/X* chiasmate homologs [45].

The increased frequency of arm cohesion defects that we observe in *mtrm^KG^/+* oocytes (Fig 3) compared to those that are wild type for *mtrm* (Fig 5) indicate that arm cohesion defects in *mtrm* heterozygotes are not restricted to heterochromatic sequences or achiasmate *X* chromosomes. The Oligopaint arm probe that we utilized recognizes a 100 kb euchromatic region approximately 1.5 Mb from the distal end of normal (non-rearranged) *X* chromosomes. The small size of the target sequence recognized by this arm probe may enhance our ability to detect cohesion defects. Partial dissociation of cohesin from meiotic chromosomes caused by increased PLK activity in *mtrm^KG^/+* oocytes during late prophase is likely responsible for the increased frequency of arm cohesion defects that we observe in *mtrm^KG^/+* (Fig 3) oocytes compared to *mtrm^+^* (Fig 5). This basal level of cohesion defects in *mtrm^KG^/+* control oocytes is further increased by knockdown of the cohesin subunit Smc3 during meiotic prophase (Fig 3). Our FISH data argue that *mtrm^KG^/+* oocytes provide a useful sensitized background for measuring segregation errors caused by premature loss of arm cohesion not only because the achiasmate system is disabled but also because this genotype reduces (but does not eliminate) the number of functional cohesive linkages on meiotic chromosomes during late prophase.

Our results indicate that turnover of cohesin and formation of new cohesive linkages during meiotic prophase are required to prevent premature loss of arm cohesion in Drosophila oocytes during normal oogenesis. These findings conflict with the popular model that meiotic cohesion in mammalian oocytes is only established before birth and that gradual loss of these original cohesive linkages during aging leads to increased segregation errors in the oocytes of older women. Although analysis of mouse oocytes has not uncovered evidence of cohesin turnover during meiotic prophase [15–17], the mammalian cohesin loader (NIPBL) and cohesion establishment factor (ESCO2) localize to mouse oocyte chromosomes during multiple prophase stages, including diplotene [46–48]. In addition, a recent report indicates that loss of ESCO2 activity during prophase in mouse spermatocytes causes cohesion defects along the sex chromosome arms [49]. Our finding that cohesin mislocalization occurs in Drosophila oocytes when the normal stoichiometry of cohesin subunits is altered (Fig S1 and S4) echoes that of a previous study using HeLa cells [32] and illustrates an inherent experimental challenge when testing the hypothesis that new cohesive linkages are formed during meiotic prophase under normal conditions.

Here we demonstrate that chromosome association of cohesin is highly dynamic during meiotic prophase and that cohesin loading and establishment of new cohesive linkages after oocyte DNA replication is required to maintain arm cohesion in Drosophila oocytes. However, even with a cohesion rejuvenation program that operates during meiotic prophase, diplotene-like Drosophila oocytes that undergo aging exhibit a significant increase in chromosome missegregation [23, 40]. Furthermore, our previous work revealed that oxidative damage contributes to age-induced meiotic segregation errors [23, 33]. Together, our findings raise the possibility that oxidative damage compromises the rejuvenation program in aging Drosophila oocytes. If a meiotic cohesion rejuvenation pathway also operates in human oocytes, premature loss of cohesion may arise in the oocytes of older women, at least in part, because rejuvenation declines with age. Further delineation of the mechanism(s) underlying rejuvenation and its possible age-induced decline could potentially inform therapeutic strategies to bolster rejuvenation and slow cohesion loss in aging oocytes.

## ACKNOWLEDGEMENTS

We thank the Bloomington Drosophila Stock Center (NIH P40OD018537), the Transgenic RNAi Project (NIH R24OD030002), and the Drosophila Genomics Resource Center (NIH 2P40OD010949) for providing flies and reagents. We thank R. Scott Hawley and Greg Rogers for providing C(3)G and CID antibodies, respectively. We are grateful to Amrita Sontakke for technical assistance, Britton Johnson for preparation of fly food, Ann Lavanway for assistance with microscopy, and Jeff Butler at Volocity for technical support. We thank Roger Sloboda, Amanda Amodeo and members of the Bickel lab for helpful comments. This work was funded by NIH R01GM05934 awarded to SEB.

## AUTHOR CONTRIBUTIONS

KAW generated constructs and flies to express HA-tagged Smc3*. MAH and SEB designed the experiments. MAH conducted the experiments and analyzed the data. MAH and SEB wrote the manuscript.

## SUPPLEMENTARY INFORMATION

### STAR Methods

**Table S1:** Primers used for site directed mutagenesis and Smc3*-HA cloning

**Table S2:** Fly stocks and genotypes

**Figure S1:** Overexpression of HA-tagged-Smc3* disrupts its normal localization.

**Figure S2:** The Smc1 tag-switch transgene produces physiological levels of tagged-Smc1 protein that is functional.

**Figure S3:** sGFP-tagged Smc1 is not detectable until after FLP expression is induced by the matα-Gal4 driver.

**Figure S4:** Matα-induced knockdown of endogenous Smc3 prevents nuclear localization of Smc1-sGFP in late prophase oocytes.

## STAR METHODS

### Creation of WT-Smc3*-HA and KK-Smc3*-HA transgenic flies

All site-directed mutagenesis was performed using the QuikChange II XL Site-Directed Mutagenesis Kit (Agilent 200521) and PAGE purified primers from Integrated DNA Technologies. All mutations were verified by sequencing. Primer sequences are provided in Table S1. Using a Bluescript KS vector containing 12 tandem HA tags (12X HA) (gift from Christian Lehner, [1]), we engineered a stop codon at the end of the HA open reading frame (ORF). The Smc3 ORF (including the consensus translational start sequence [2]) was PCR amplified using clone RE14758 (DGRC Stock 8502; https://dgrc.bio.indiana.edu//stock/8502; RRID:DGRC_8502) obtained from the Drosophila Genomics Resource Center. PCR primers introduced Kpn1 and Not1 sites at the 5’ end and an Xho1 site at the 3’ end to facilitate subcloning. Following Kpn1 and Xho1 digestion, the amplified Smc3 fragment was cloned into the above KS vector, immediately upstream of the 12X HA sequence. The entire ORF was sequenced to verify that no mutations occurred during PCR amplification.

Silent mutations were introduced into the Smc3 sequence so that the encoded mRNA (Smc3*) was insensitive to the Smc3 short hairpin SH00137.N used to knock down endogenous Smc3. The original Smc3 sequence: GAGTATATACGCTACGAA was mutated to GA<u>A</u>TA<u>C</u>AT<u>T</u>CG<u>G</u>TA<u>T</u>GAG (mutated nucleotides underlined), resulting in WT-Smc3*-HA.

Eco-dependent acetylation of two conserved lysines is required for cohesion establishment during S phase [3–7]. For the KK-Smc3*-HA construct, site-directed mutagenesis was used to convert these lysines (K105, K106 in Drosophila Smc3) to arginines, which cannot be acetylated. WT-Smc3*-HA and KK-Smc3*-HA were each subcloned into pUASP-attB [8] using Not1 and Spe1 sites and confirmed by sequencing. Injections were performed by Genetic Services, Inc (Sudbury, MA) to generate PhiC31 integrase mediated *attP40* insertions on chromosome 2 [9].

### Generation of Smc1 tag-switch construct and transgenic flies

The Smc1 tag-switch construct (see Fig 4) was synthesized by Genscript Biotech Corp (Piscataway, NJ) using the Drosophila *smc1* genomic sequence (FlyBase ID FBgn0040283). In addition to all exons and introns, 976 bp of upstream regulatory DNA (including the promoter) as well as the 5’ and 3’UTRs were included. At the end of the Smc1 ORF, a 48bp FRT site replaces the Smc1 stop codon and is followed by the mCherry coding sequence, a 3X HA tag (with stop codon), the Smc1 3’ UTR, and an additional 100bp of *smc1* genomic DNA. Immediately downstream of the mCherry cassette lies an additional 48 bp FRT site, followed by DNA encoding superfolder GFP (sGFP), an EPEA epitope tag (with stop codon), the *smc1* 3’ UTR, and an additional 100bp of *smc1* genomic DNA. Following synthesis, GenScript cloned the construct into the pUC57 vector, verified the sequence, and subcloned the insertion into the w+attB vector (gift from Jeff Sekelsky, Addgene plasmid # 30326; http://n2t.net/addgene:30326). Both mCherry and sGFP sequences were codon optimized for Drosophila. Sequence of the Smc1 tag-switch construct in the w+attB vector will be provided upon request. BestGene Inc (Chino Hills, CA) performed injections to insert the Smc1 tag-switch transgene from w+attB into the *attP40* landing site on chromosome 2 [9].

### Fly stocks and crosses

All fly stocks and crosses were reared on standard cornmeal molasses medium in a humidified chamber at 25° C. Table S2 provides full genotypes for stocks used in this study, and Bickel stock numbers from this table are noted below for cross descriptions.

To induce expression of UASp-Smc3*-HA, we utilized two previously described [10] matα-Gal4-VP16 driver chromosomes (“strong” and “weak”), each of which also contains the *mtrm^KG08051^* allele (W-110 and W-107). Expression induced by our strong matα driver is similar to that of the original matα chromosome (Bloomington #7063, T-273). Gal4-inducible expression is considerably lower when we utilize our weak matα driver chromosome [10].

To knock down endogenous Smc3 during meiotic prophase and simultaneously express wild-type Smc3*-HA (WT), mutant Smc3*-HA (KK) or no additional Smc3 protein (Empty), *strong matα, mtrm^KG^ /TM3 (*W-110) *or weak matα, mtrm^KG^ /TM3* (W-107) males were crossed to *UAS-WT-Smc3*-HA; Smc3 hairpin* (T-622), *UAS-KK-Smc3*-HA; Smc3 hairpin* (T-625) or *UAS-Empty; Smc3 hairpin* (T-800) virgins. Non-balancer female progeny were used for immunostaining. The strong matα driver was used for immunolocalization of HA-tagged WT– and KK-Smc3* (Figs 1C, S1) as well as quantification of SC defects (Fig 2). The weak matα driver was also used to assay SC defects (Fig 2) and to quantify cohesion defects using FISH (Fig 3).

For Smc1 tag-switch experiments, we used the original matα-Gal4-VP16 driver chromosome (T-273, *mtrm^+^*) to induce expression of Flippase (FLP) after meiotic S phase. Excision of FRT-flanked mCherry sequences in the Smc1 tag-switch transgene induces a switch from Smc1-mCherry to Smc1-GFP expression during meiotic prophase. *UASp-FLP; matα* (T-778) males were crossed to *Smc1 tag-switch* (T-772) or *Smc1 tag-switch; Smc3 hairpin* (T-779) virgins. Ovaries of female progeny were used for immunolocalization of Smc1-mCherry, Smc1-sGFP, and CID on meiotic chromosomes (Figs 4, 5, S3, S4) and for FISH analysis of cohesion defects (Fig 5).

To confirm that Smc1-mCherry and Smc1-sGFP were each functional, we performed a series of crosses to determine whether the lethality of flies transheterozygous for two different *smc1* deletion alleles (*smc1Δ^ex46^*/*smc1Δ^ex89^*) was rescued when the Smc1 tag-switch transgene was present and solely expressing either Smc1-mCherry or Smc1-sGFP. In the absence of FLP, the intact Smc1 tag-switch transgene expresses only mCherry-tagged Smc1. Following a series of crosses (using T-815, OL-124 and M-743), we recovered S*mc1-mCherry/+; smc1Δ^ex46^/smc1Δ^ex89^*males and females at the expected ratio and verified that they were fully fertile (Fig. S2B). To generate flies solely expressing Smc1-sGFP, we generated a stock (T-816, *Smc1-sGFP*) starting with progeny from *UASp-FLP/+; Smc1 tag-switch*/+; *matα/+* females. Progeny that inherit the FRT-excised transgene (Smc1-sGFP) can no longer express Smc1-mCherry. We confirmed the absence of Smc1-mCherry expression in this stock using immunostaining. Through a series of crosses (using T-817, OL-124 and M-743), we verified that S*mc1-sGFP/+; smc1Δ^ex46^/smc1Δ^ex89^*males and females eclosed at the expected ratio and were fully fertile (Fig. S2B).

### Immunostaining

Newly eclosed females were fattened for 2-3 days in vials containing food, yeast, and males. Following dissection in 1X PBS, ovarioles were partially separated with a fine tungsten needle and fixed for 20 minutes in a mixture of 600µl heptane and 200µl of 2% formaldehyde (Ted Pella, 18505) containing 0.5% NP-40 (Thermo Fisher 28324). All incubations and washes were performed in glass dishes at room temperature (RT) with gentle agitation on a rotating platform unless otherwise stated. Fixed ovaries were rinsed three times in 1X PBST (1X PBS + 0.2% Tween-20, Thermo Fisher 28320), blocked in 1X PBST + 1% BSA (Fisher BP1605-100) for 1 hour, and incubated at 4°C overnight in antibody incubation buffer (1X PBS + 0.01% Tween-20 + 1% BSA) containing the appropriate primary antibodies. After rinsing three times in 1X PBST, ovaries were washed three times for 20 minutes in 1X PBST and placed in antibody incubation buffer containing the appropriate secondary antibodies for 1 hour. Following three rinses in 1X PBST, one 20-minute wash in 1X PBST, a 20-minute incubation in 1µg/ml DAPI in 1X PBS, and one 20-minute wash in 1XPBS + 0.01% Tween-20, ovarioles were fully separated with tungsten needles. Ovarioles were transferred onto to 18mm poly-L-lysine coated #1.5 coverslips, and excess liquid removed before adding mounting media.

For HA-tagged-Smc3* and C(3)G immunolocalization (Figures 1C, 2A and S1) and quantification of SC defects (Figure 2), high-affinity rat anti-HA (Roche 3F10, 1:2000) and mouse anti-C3G (1A8-1G2, 1:1000, gift from Hawley lab [11]) were used. Cy3 anti-rat (min-X mouse, 712-165-153) and Alexa 488 anti-mouse (min-X rat, 715-545-151) secondary antibodies from Jackson ImmunoResearch were used at 1:400. Slides were mounted in SlowFade Diamond Antifade (Molecular Probes S36967, Figs 1 & 2) or Prolong Gold Antifade (Molecular Probes P36930, Figs 2 & S1).

For Smc1-mCherry and Smc1-sGFP detection (Figures 4, 5, S3 and S4), GFP-Booster Alexa Fluor 488 (Alpaca anti-GFP, ChromTek gb2AF488) and RFP-Booster Alexa Fluor 568 (Alpaca anti-mCherry, ChromTek rb2AF568) primary antibodies were each used at 1:500 (with no secondary antibody). To visualize centromeres, rabbit anti-CID (1:10,000, gift from Rogers lab, [12]) was used followed by Cy5 anti-rabbit (Jackson ImmunoResearch 711-175-152, 1:400). Samples were incubated overnight at 4°C with primary antibodies (no rotation) and 1 hr at RT with the secondary antibody. Slides were mounted in Vectashield Vibrance (Vector Laboratories, H-1700, Fig 4) or SlowFade Diamond (Figs 5, S3 & S4)

## FISH

For detecting arm cohesion defects in mature Drosophila oocytes (stage 13-14), we utilized an Alexa 647-labeled Oligopaint probe (OPP122) generated by the Joyce Lab, University of Pennsylvania. This mixture of oligonucleotides (each with 80 bases of homology) contains 937 unique oligos targeting a 100kb distal region of the X chromosome (dm6, nucleotides 1,400,000-1,500,000) and was used at a final concentration of 0.50pmol/µl. Cy3-conjugated probe (5′-Cy3-AGGGATCGTTAGCACTCGTAAT; Integrated DNA Technologies) that hybridizes to the 359-bp repeat in pericentric heterochromatin of the X chromosome was used at a concentration of 1ng/µl. Fixation, hybridization, and washes were performed as previously described [13, 14], except that oocytes were subjected to a pre-denaturation step (5min @ 37°C, 3 min @ 92°C, 20 min @ 60°C, hold @ 37°C) prior to denaturation and hybridization (5min @ 37°C, 3 min @ 92°C, hold @ 37°C overnight). Oocytes were mounted on 18 mm #1.5 coverslips in Prolong Gold mounting medium and slides were allowed to cure in a box containing desiccant for at least 14 days in the dark.

### Image acquisition, processing, and quantitative analysis

All images were acquired using an Andor Spinning Disk confocal on a Nikon Eclipse Ti inverted microscope equipped with an ASI MS-2000 motorized piezo stage and Zyla 4.2-megapixel sCMOS camera. Image acquisition utilized Nikon Elements software (5.11.02 Build 1369), a 50 µm pinhole disk, and up to four laser lines (405, 488, 561 and 637 nm). A Nikon CFI 60X Plan Apo oil objective (NA 1.4) was utilized for immunostaining experiments, and a Nikon CFI 100X oil Plan Apo DIC objective (NA 1.45) was used for FISH experiments. All image acquisition utilized 4X frame averaging. Z-series acquisition was as follows: Smc3*-HA (0.3 µm steps, 3 µm total), C(G)3 in Fig 2 (0.1 µm steps, 6 µm total), Smc1 tag-switch (0.5 µm steps, 2 µm total), FISH (0.1 µm steps, 6 µm total). For multi-channel Z-series, an entire Z-stack was captured for one fluor before switching to the next laser. When imaging oocyte chromatin in intact ovarioles, only a specified ROI was captured, but we also collected a full-frame single-plane DAPI image of the ovariole using a Nikon CFI 20X Plan Apo oil objective (NA 0.75) which allowed us to stage individual egg chambers using size and morphological criteria [15–17]. For all immunostaining comparisons presented in figures, identical acquisition, deconvolution and processing were used for different genotypes, including the number of optical sections included in the projection shown.

#### Scoring for SC defects

To quantify SC defects, ovarioles stained with anti-C(3)G antibody were viewed using a 63X Plan-Apochromat (NA 1.4) objective on a Zeiss AxioImager M1 microscope equipped with a Hamamatsu ORCA^R2^ digital camera controlled by Volocity Acquisition software (v6.5.1). Slides were blinded, and scoring was performed by viewing an enlarged C(3)G image on the computer monitor while slowly focusing up and down through the oocyte nucleus. Oocytes were analyzed from germarial region 3 through stage 6, and the SC was determined to be normal (continuous, long threads) or to exhibit premature disassembly (broken, short, or spots indicate increasing severity of SC defects).

#### Quantification of Smc1-mCherry and Smc1-sGFP on oocyte chromosomes

Volocity Quantification (v6.5.0) was used to quantify Smc1-sGFP and Smc1-mCherry signal intensities on non-deconvolved confocal Z series. For germarial region 3 to stage 6, the volume occupied by oocyte DNA (405 channel) was cropped using a free-hand tool while viewing a maximum intensity projection (5 steps, 2 µm total). Smc1 signal in the cropped volume was quantified separately for mCherry and sGFP using a threshold corresponding to 0.5 standard deviation (SD). For each volume, centromeres were identified using CID intensity thresholding (637 nm channel, 3.0 SD). For each oocyte, total intensity was calculated for the centromere volume and this value was subtracted from the total Smc1-mCherry or Smc1-sGFP signal on the DNA volume to determine the total signal intensity on chromosome arms. Average intensity per voxel was calculated and graphed for arms and centromeres. For stage 10 oocytes, voxels containing oocyte DNA were identified by thresholding the DAPI signal (0.5 SD) and the Smc1-sGFP signal on DNA was quantified.

#### Scoring cohesion defects

All FISH scoring was performed blind to genotype. Following image acquisition, a MATLAB (R2022a) script was generated and used to randomly mix and rename Nikon Elements. nd2 image stacks from two or more genotypes and output a spreadsheet with the key. Blinded Z-stacks were deconvolved (Volocity Restoration v6.5.0) and visualized in three dimensions (Volocity Visualization v6.5.0) to determine the number of X chromosome arm (Alexa 647) and pericentric (Cy3) spots present on the oocyte DNA. Probe signals were considered separate if they were separated in all three dimensions by at least half the diameter of the smallest spot. 3D scoring was essential to rule out connections in the Z dimension. Two spots connected by a thread-like signal were common and were not scored as separate foci. Following scoring, the MATLAB generated key was used to determine the genotype associated with each image stack, and the percentage of oocytes with cohesion defects was calculated. A two-tailed Fisher’s exact test (Graphpad) was used to determine whether the incidence of defects in two genotypes was statistically significant (P < 0.05).

### Protein extracts and immunoblotting

Young females were fattened for 3 days in food vials with yeast and males. 20 sets of ovaries were dissected in 1X PBS, flash frozen in a microtube, and stored at –80°C. Frozen ovaries were homogenized in 200µl RIPA buffer (Sigma-Aldrich R0278) with 1X HALT protease inhibitor (Thermo Fisher 87786) using a motorized disposable pestle for 30 seconds. Benzonase digestion (Millipore Sigma E1014, 250 units, 15 min) on a nutator at RT was stopped by addition of EDTA to 5mM (Thermo Fisher 1861274). The sample was filtered through a 0.22µm centrifugal filter (Millipore UFC30GV00) at 12,000 x g for 4 minutes at RT and stored in aliquots at –80°C.

Total ovary extract was separated by SDS/PAGE on a 7.5% protein gel (BioRad Stain-Free) and transferred to Immobilon-P membrane (Millipore IPVH00010). After blocking overnight in 1X TBS containing 2% BSA and 5% non-fat dry milk, blots were incubated for one hour at RT in antibody incubation buffer (1X TBS + 0.1% Tween-20 [BioRad 1610781] + 5% non-fat dry milk) with a 1:1000 dilution of affinity-purified guinea pig anti-Smc1 [18] or rabbit anti-GFP (Invitrogen A11122). Following washes, blots were incubated for 30 min in alkaline phosphatase conjugated secondary antibody (1:3000 AP goat anti-guinea pig [Southern Biotech 6090-04] or 1:7500 goat anti rabbit [Promega S373B]) and washed before applying substrate (BioRad 1705018) for five minutes. Blots were imaged using a ChemiDoc Touch system (BioRad), and bands were quantified using BioRad Image Lab software (Version 6.1).

**Table S1:**
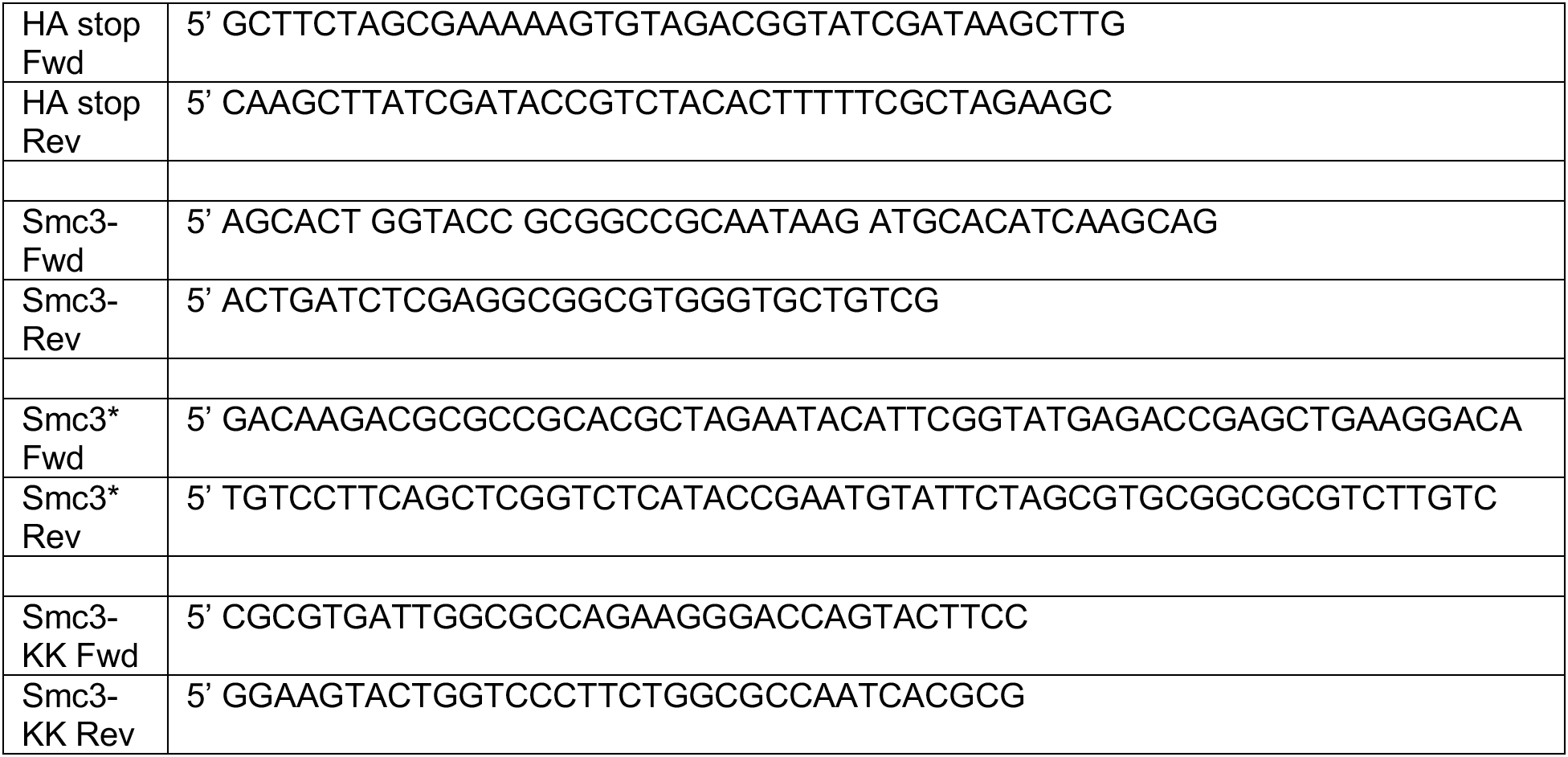
Primers used for site-directed mutagenesis and Smc3*-HA cloning.

**Table S2:**
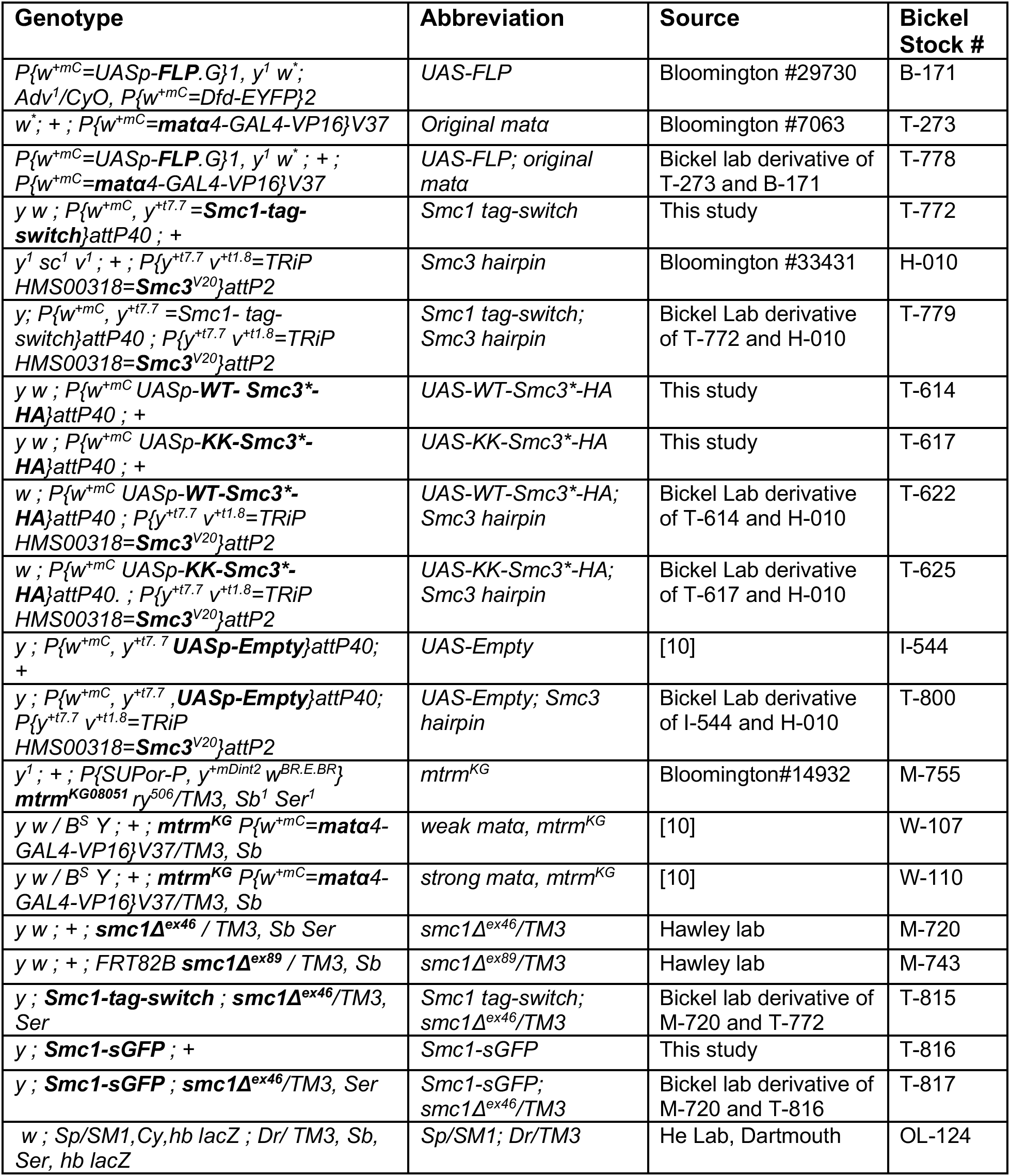
Fly Stocks and Genotypes. Smc3 hairpin: SH00137.N hairpin in Valium 20 inserted at the attP2 site (FBti0140240)

---

**Figure S1:**
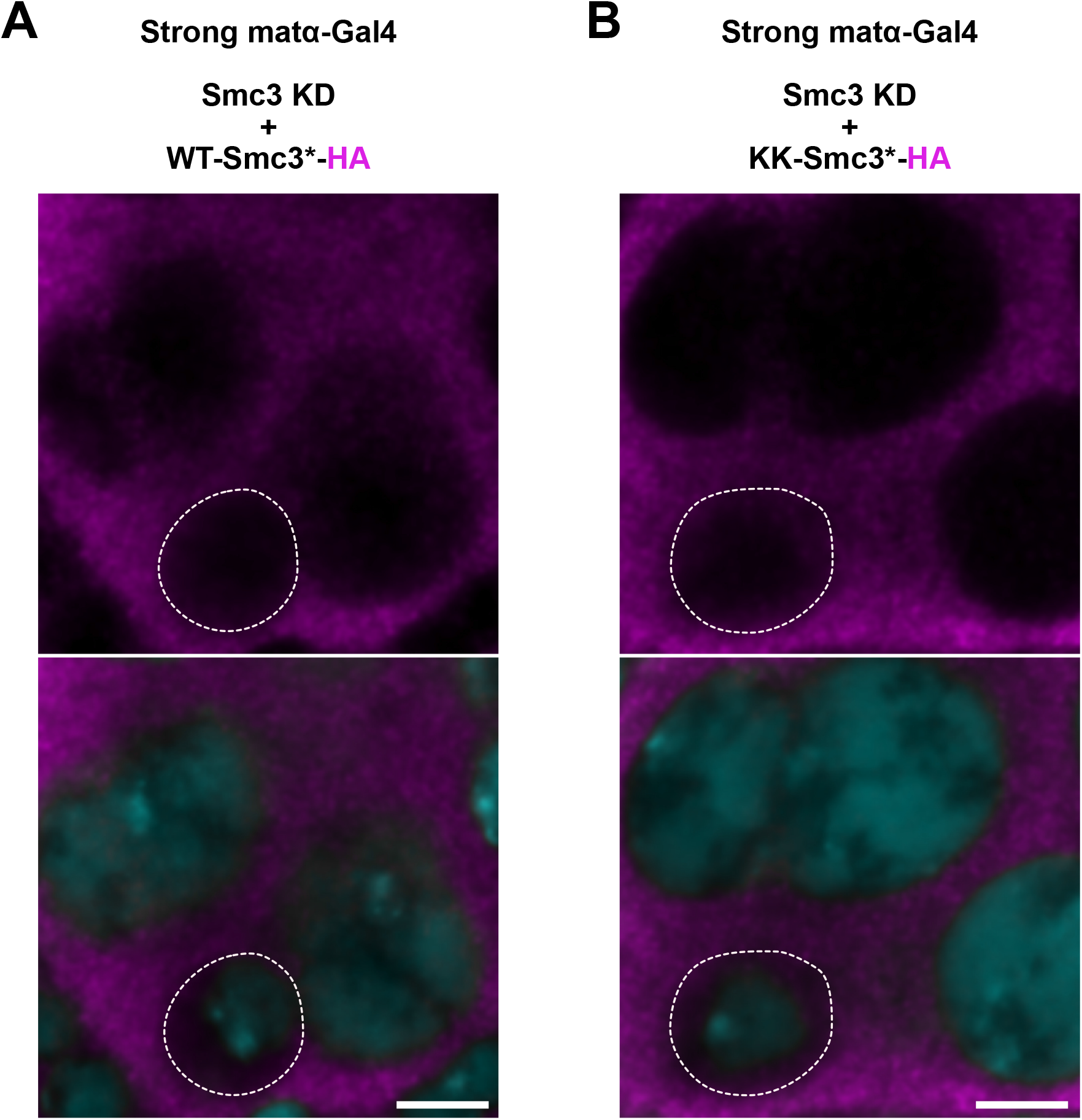
Overexpression of HA-tagged-Smc3* disrupts its normal localization. Single optical sections of deconvolved confocal Z series are shown for Smc3*-HA localization (magenta) and DNA (cyan) in stage 5 egg chambers in which expression of the Smc3 hairpin and Smc3*-HA are induced by the strong matα-Gal4 driver. Dotted lines demark the perimeter of the oocyte nucleus. In contrast to earlier stages in which Smc3*-HA localizes to oocyte chromosomes (see Fig 1C), HA-tagged Smc3* in later stages is found predominantly in the cytoplasm with little to no signal in nuclei. Because matα-Gal4 induced expression increases substantially during oogenesis [19], mislocalization of Smc3*-HA is likely due to its overexpression in later stages. Scale bar, 5µm.

**Figure S2:**
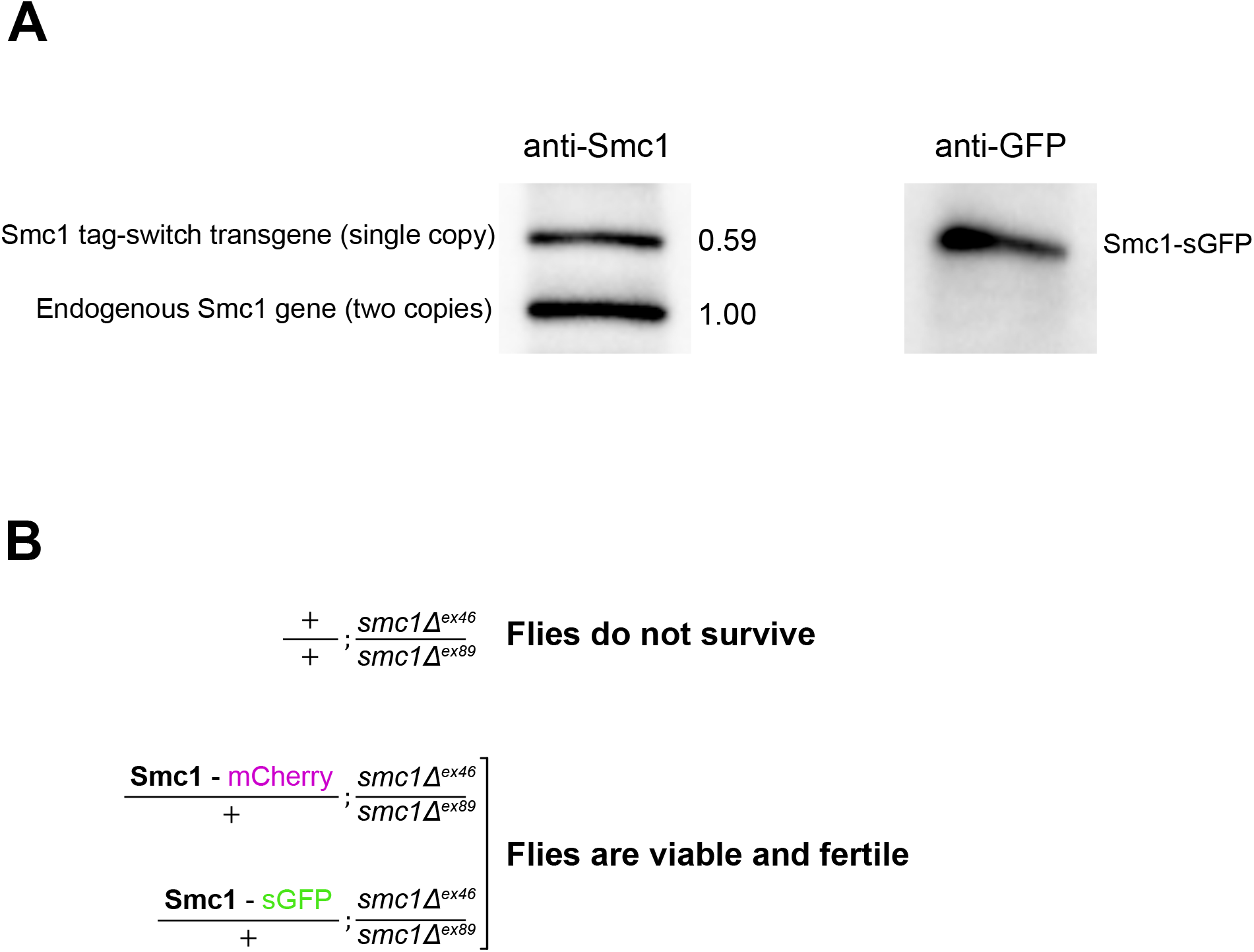
The Smc1 tag-switch transgene produces physiological levels of tagged-Smc1 protein that is functional. **A**. The indicated antibodies were used to detect Smc1 protein in total ovary extract prepared from females containing a single Smc1 tag-switch transgene. In these flies, UAS-FLP expression induced by the original matα-Gal4 driver chromosome elicits a switch from Smc1-mCherry to Smc1-sGFP expression during meiotic prophase. Two lanes from the same blot were cut and processed individually. The anti-GFP blot confirms that the upper band on the anti-Smc1 blot corresponds to Smc1 expressed from the tag-switch transgene. Based on quantification of anti-Smc1 band intensities (4 independent blots), Smc1 protein expressed from a single copy of the transgene corresponds to 59% of the signal for untagged-Smc1 (expressed from the two endogenous *smc1* alleles). **B.** Presence of a single copy of the Smc1 tag-switch transgene rescues the lethality caused by deletion of both copies of the endogenous *smc1* gene. *smc1Δ^ex46^/ smc1Δ^ex89^* transheterozygotes expressing solely mCherry-tagged Smc1 or sGFP-tagged Smc1 were fully viable and fertile.

**Figure S3.**
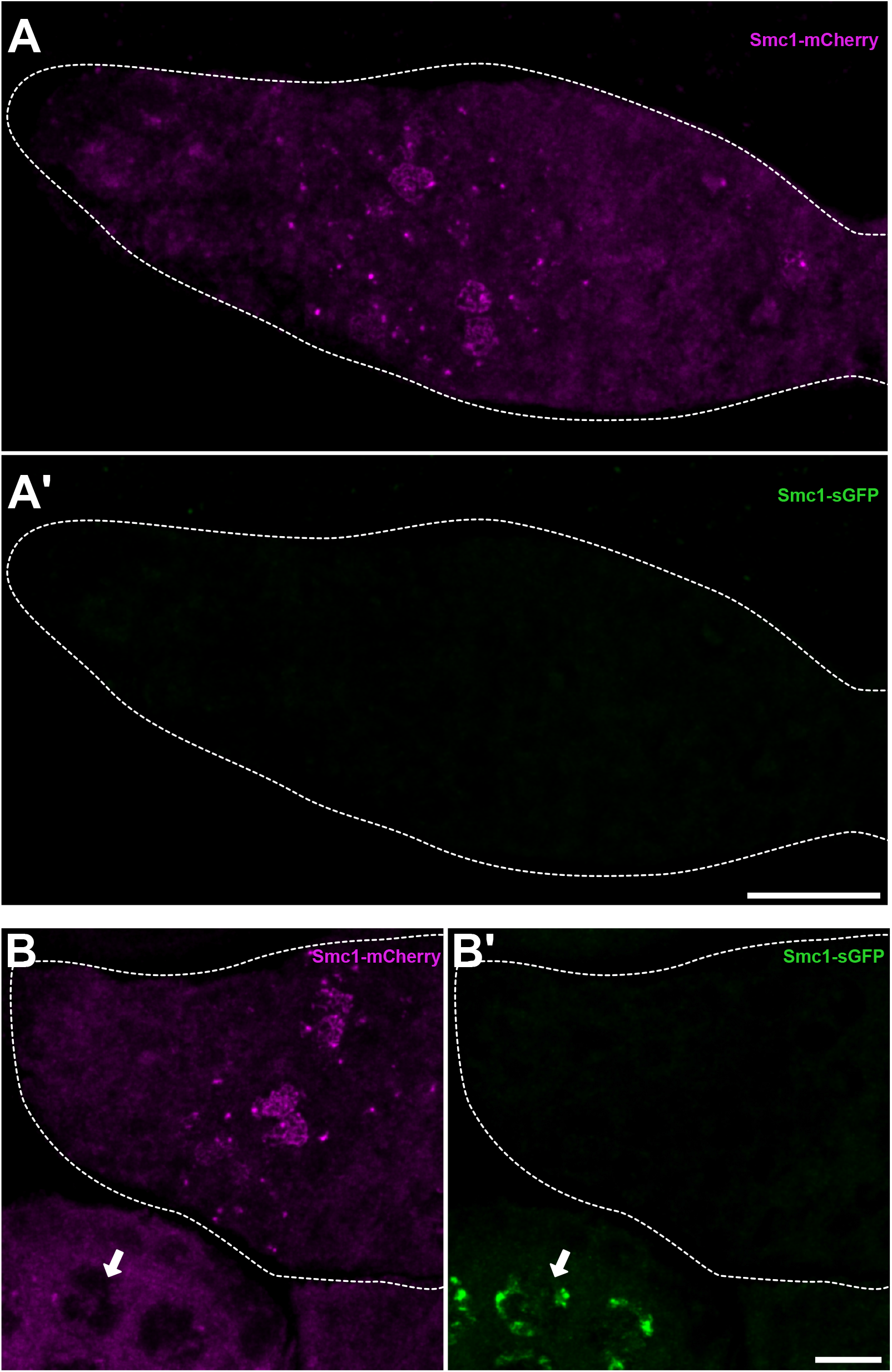
sGFP-tagged Smc1 is not detectable until after FLP expression is induced by the matα-Gal4 driver. **A and A’**. In females in which the matα-Gal4 driver induces germline expression of a UAS-FLP transgene, only Smc1-mCherry (magenta) is visible within the germarium. Expression of Smc1-sGFP (green) from the Smc1 tag-switch transgene is not detectable before the matα-Gal4 driver turns on. **B and B’.** A second germarium is shown for the same genotype. Although Smc1-sGFP signal is absent within the germarium, chromosome association of sGFP-tagged Smc1 is visible within nurse cell nuclei of a nearby stage 4 egg chamber. Arrow points to a single nurse cell nucleus. Images are maximum intensity projections of confocal Z series with a dotted line marking the perimeter of the germarium and the anterior on the left. Scale bars, 15 µm. Genotype is the same as that indicated in Fig 4B.

**Figure S4:**
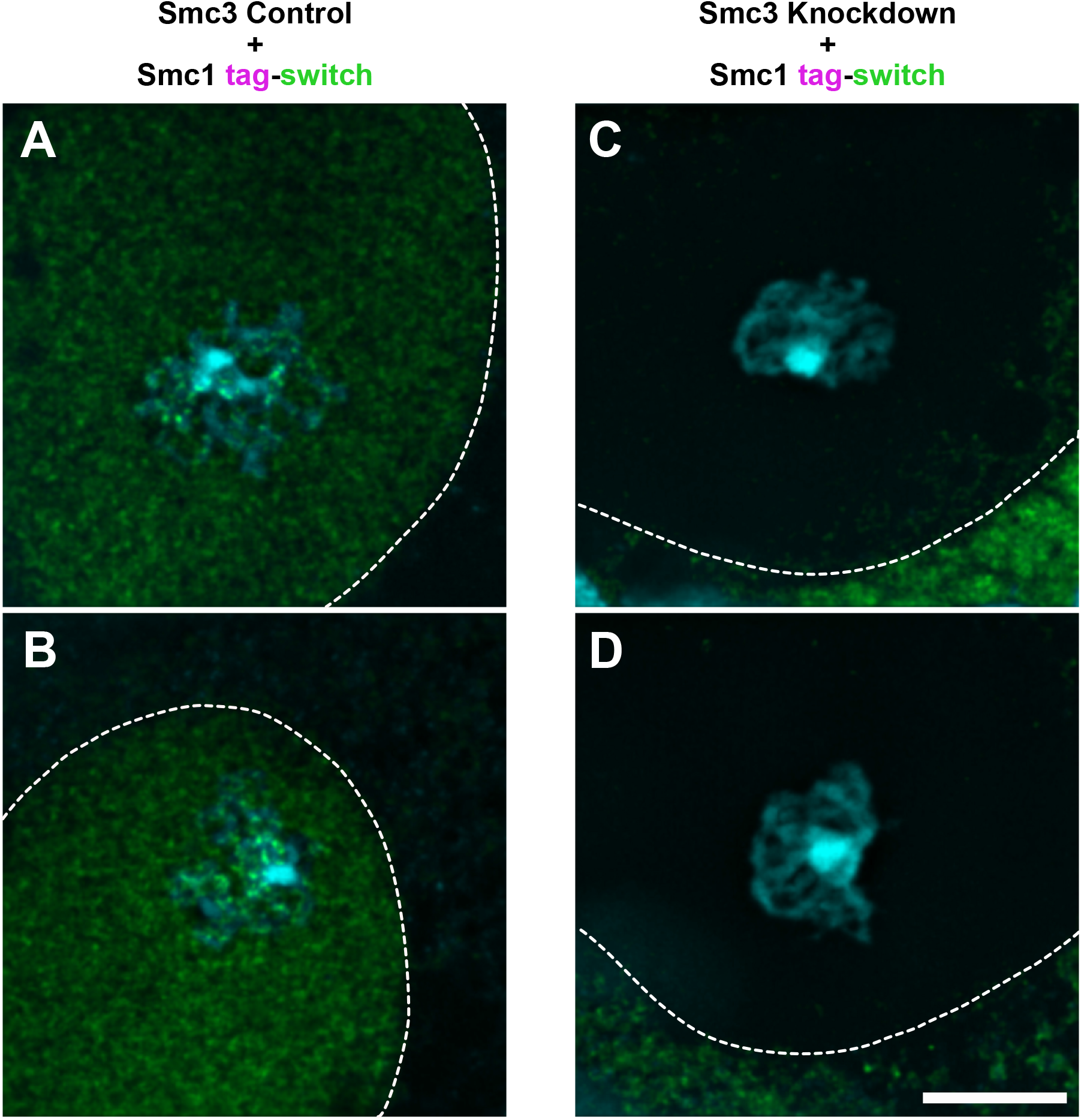
Matα-Gal4 induced knockdown of endogenous Smc3 prevents nuclear localization of Smc1-sGFP in late prophase oocytes. DNA (cyan) and sGFP-tagged Smc1 (green) within stage 10 egg chambers are shown for two different genotypes. Only a portion of the oocyte nucleus is visible, with a dotted line marking the nuclear/cytoplasmic boundary. Images are maximum intensity projections of deconvolved confocal Z series. Scale bar, 8µm. **A & B.** When endogenous Smc3 levels are not perturbed (Smc3 Control) and expression of matα-Gal4 drives FLP-induced expression of Smc1-sGFP, localization of sGFP-tagged Smc1 is enriched in the oocyte nucleus and associates with the meiotic chromosomes. **C & D**. When the matα-Gal4 driver induces expression of FLP as well as the Smc3 hairpin (Smc3 Knockdown), sGFP-tagged Smc1 is excluded from the oocyte nucleus and fails to localize to the meiotic chromosomes. Note that the top and bottom panels in Fig 5A correspond to cropped versions of A and C, respectively. Genotypes are identical to those indicated in Fig 4B (Smc3 Control) and Fig 4C (Smc3 Knockdown).

## REFERENCES CITED

1. Nasmyth, K. and C.H. Haering, Cohesin: its roles and mechanisms. Annu Rev Genet, 2009. 43: p. 525–58.

2. Peters, J.M. and T. Nishiyama, Sister chromatid cohesion. Cold Spring Harbor Perspectives in Biology, 2012. 4(11).

3. McNicoll, F., M. Stevense, and R. Jessberger, Cohesin in gametogenesis. Current Topics in Developmental Biology, 2013. 102: p. 1–34.

4. Marston, A.L., Chromosome segregation in budding yeast: sister chromatid cohesion and related mechanisms. Genetics, 2014. 196(1): p. 31–63.

5. Morales, C. and A. Losada, Establishing and dissolving cohesion during the vertebrate cell cycle. Curr Opin Cell Biol, 2018. 52: p. 51–57.

6. Ishiguro, K.I., The cohesin complex in mammalian meiosis. Genes Cells, 2019. 24(1): p. 6–30.

7. Ivanov, D., et al., Eco1 Is a Novel Acetyltransferase that Can Acetylate Proteins Involved in Cohesion. Curr. Biol., 2002. 12(4): p. 323–8.

8. Ben-Shahar, T.R., et al., Eco1-dependent cohesin acetylation during establishment of sister chromatid cohesion. Science, 2008. 321(5888): p. 563–6.

9. Unal, E., et al., A molecular determinant for the establishment of sister chromatid cohesion. Science, 2008. 321(5888): p. 566–9.

10. Zhang, J., et al., Acetylation of Smc3 by Eco1 is required for S phase sister chromatid cohesion in both human and yeast. Mol Cell, 2008. 31(1): p. 143–51.

11. Song, J., et al., Cohesin acetylation promotes sister chromatid cohesion only in association with the replication machinery. The Journal of biological chemistry, 2012. 287(41): p. 34325–36.

12. Greaney, J., Z. Wei, and H. Homer, Regulation of chromosome segregation in oocytes and the cellular basis for female meiotic errors. Hum Reprod Update, 2018. 24(2): p. 135–161.

13. Wartosch, L., et al., Origins and mechanisms leading to aneuploidy in human eggs. Prenat Diagn, 2021. 41(5): p. 620–630.

14. Charalambous, C., A. Webster, and M. Schuh, Aneuploidy in mammalian oocytes and the impact of maternal ageing. Nat Rev Mol Cell Biol, 2023. 24(1): p. 27–44.

15. Revenkova, E., et al., Oocyte cohesin expression restricted to predictyate stages provides full fertility and prevents aneuploidy. Curr Biol, 2010. 20(17): p. 1529–33.

16. Tachibana-Konwalski, K., et al., Rec8-containing cohesin maintains bivalents without turnover during the growing phase of mouse oocytes. Genes Dev, 2010. 24(22): p. 2505–16.

17. Burkhardt, S., et al., Chromosome Cohesion Established by Rec8-Cohesin in Fetal Oocytes Is Maintained without Detectable Turnover in Oocytes Arrested for Months in Mice. Curr Biol, 2016. 26(5): p. 678–85.

18. Jessberger, R., Age-related aneuploidy through cohesion exhaustion. EMBO reports, 2012. 13(6): p. 539–46.

19. Weng, K.A., C.A. Jeffreys, and S.E. Bickel, Rejuvenation of Meiotic Cohesion in Oocytes during Prophase I Is Required for Chiasma Maintenance and Accurate Chromosome Segregation. PLoS Genetics, 2014. 10(9): p. e1004607.

20. Strom, L., et al., Postreplicative recruitment of cohesin to double-strand breaks is required for DNA repair. Molecular Cell, 2004. 16(6): p. 1003–15.

21. Strom, L., et al., Postreplicative formation of cohesion is required for repair and induced by a single DNA break. Science, 2007. 317(5835): p. 242–5.

22. Unal, E., J.M. Heidinger-Pauli, and D. Koshland, DNA double-strand breaks trigger genome-wide sister-chromatid cohesion through Eco1 (Ctf7). Science, 2007. 317(5835): p. 245–8.

23. Perkins, A.T., et al., Increased levels of superoxide dismutase suppress meiotic segregation errors in aging oocytes. Chromosoma, 2019. 128(3): p. 215–222.

24. Harris, D., et al., A deficiency screen of the major autosomes identifies a gene (matrimony) that is haplo-insufficient for achiasmate segregation in Drosophila oocytes. Genetics, 2003. 165(2): p. 637–52.

25. Cahoon, C.K. and R.S. Hawley, Regulating the construction and demolition of the synaptonemal complex. Nat Struct Mol Biol, 2016. 23(5): p. 369–77.

26. Ur, S.N. and K.D. Corbett, Architecture and Dynamics of Meiotic Chromosomes. Annu Rev Genet, 2021. 55: p. 497–526.

27. Adams, I.R. and O.R. Davies, Meiotic Chromosome Structure, the Synaptonemal Complex, and Infertility. Annu Rev Genomics Hum Genet, 2023.

28. Llano, E. and A.M. Pendas, Synaptonemal Complex in Human Biology and Disease. Cells, 2023. 12(13).

29. Khetani, R.S. and S.E. Bickel, Regulation of meiotic cohesion and chromosome core morphogenesis during pachytene in Drosophila oocytes. J Cell Sci, 2007. 120(Pt 17): p. 3123–37.

30. Page, S.L. and R.S. Hawley, *c(3)G encodes a Drosophila synaptonemal complex protein*. Genes Dev, 2001. 15(23): p. 3130–43.

31. Wesley, E.R., R.S. Hawley, and K.K. Billmyre, Genetic background impacts the timing of synaptonemal complex breakdown in Drosophila melanogaster. Chromosoma, 2020. 129(3-4): p. 243–254.

32. Laugsch, M., et al., Imbalance of SMC1 and SMC3 Cohesins Causes Specific and Distinct Effects. PloS One, 2013. 8(6): p. e65149.

33. Perkins, A.T., et al., Oxidative stress in oocytes during midprophase induces premature loss of cohesion and chromosome segregation errors. Proc Natl Acad Sci U S A, 2016. 113(44): p. E6823–E6830.

34. Pedelacq, J.D., et al., Engineering and characterization of a superfolder green fluorescent protein. Nat Biotechnol, 2005. 24(1): p. 79–88.

35. David, J. and J. Merle, A reevaluation of the duration of egg chamber stages in oogenesis of Drosophila melanogaster. Drosophila Information Service, 1968. 43: p. 122–123.

36. Heidinger-Pauli, J.M., et al., The kleisin subunit of cohesin dictates damage-induced cohesion. Mol Cell, 2008. 31(1): p. 47–56.

37. Heidinger-Pauli, J.M., E. Unal, and D. Koshland, Distinct targets of the Eco1 acetyltransferase modulate cohesion in S phase and in response to DNA damage. Mol Cell, 2009. 34(3): p. 311–21.

38. Hawley, R.S., et al., There are two mechanisms of achiasmate segregation in Drosophila, one of which requires heterochromatic homology. Dev. Genet., 1992. 13: p. 440–467.

39. Karpen, G.H., M.H. Le, and H. Le, Centric heterochromatin and the efficiency of achiasmate disjunction in Drosophila female meiosis. Science, 1996. 273: p. 118–122.

40. Subramanian, V.V. and S.E. Bickel, Aging predisposes oocytes to meiotic nondisjunction when the cohesin subunit SMC1 is reduced. PLoS Genet, 2008. 4(11): p. e1000263.

41. Xiang, Y., et al., The inhibition of polo kinase by matrimony maintains G2 arrest in the meiotic cell cycle. PLoS Biol, 2007. 5(12): p. e323.

42. Hauf, S., et al., Dissociation of cohesin from chromosome arms and loss of arm cohesion during early mitosis depends on phosphorylation of SA2. PLoS Biol, 2005. 3(3): p. e69.

43. Zhang, N., et al., Interaction of Sororin protein with polo-like kinase 1 mediates resolution of chromosomal arm cohesion. J Biol Chem, 2011. 286(48): p. 41826–41837.

44. Challa, K., et al., Meiosis-specific prophase-like pathway controls cleavage-independent release of cohesin by Wapl phosphorylation. PLoS Genet, 2019. 15(1): p. e1007851.

45. Bonner, A.M., S.E. Hughes, and R.S. Hawley, Regulation of Polo Kinase by Matrimony Is Required for Cohesin Maintenance during Drosophila melanogaster Female Meiosis. Curr Biol, 2020. 30(4): p. 715–722 e3.

46. Evans, E.B., et al., Localization and regulation of murine Esco2 during male and female meiosis. Biology of reproduction, 2012. 87(3): p. 61.

47. Visnes, T., et al., Localisation of the SMC loading complex Nipbl/Mau2 during mammalian meiotic prophase I. Chromosoma, 2013. 123(3): p. 239–52.

48. Kuleszewicz, K., X. Fu, and N.R. Kudo, Cohesin loading factor Nipbl localizes to chromosome axes during mammalian meiotic prophase. Cell Division, 2013. 8(1): p. 12.

49. McNicoll, F., et al., Meiotic sex chromosome cohesion and autosomal synapsis are supported by Esco2. Life Sci Alliance, 2020. 3(3).

## REFERENCES CITED

1. Heidmann, D., et al., The Drosophila meiotic kleisin C(2)M functions before the meiotic divisions. Chromosoma, 2004. 113(4): p. 177–87.

2. Cavener, D., Comparison of the consensus sequence flanking translational start sites in Drosophila and vertebrates. Nuc. Acids Res., 1987. 15: p. 1353–1361.

3. Unal, E., et al., A molecular determinant for the establishment of sister chromatid cohesion. Science, 2008. 321(5888): p. 566–9.

4. Ben-Shahar, T.R., et al., Eco1-dependent cohesin acetylation during establishment of sister chromatid cohesion. Science, 2008. 321(5888): p. 563–6.

5. Song, J., et al., Cohesin acetylation promotes sister chromatid cohesion only in association with the replication machinery. The Journal of biological chemistry, 2012. 287(41): p. 34325–36.

6. Ivanov, D., et al., Eco1 Is a Novel Acetyltransferase that Can Acetylate Proteins Involved in Cohesion. Curr. Biol., 2002. 12(4): p. 323–8.

7. Zhang, J., et al., Acetylation of Smc3 by Eco1 is required for S phase sister chromatid cohesion in both human and yeast. Mol Cell, 2008. 31(1): p. 143–51.

8. Takeo, S., et al., Shaggy/glycogen synthase kinase 3beta and phosphorylation of Sarah/regulator of calcineurin are essential for completion of Drosophila female meiosis. Proceedings of the National Academy of Sciences of the United States of America, 2012. 109(17): p. 6382–9.

9. Markstein, M., et al., Exploiting position effects and the gypsy retrovirus insulator to engineer precisely expressed transgenes. Nature genetics, 2008. 40(4): p. 476–83.

10. Perkins, A.T., et al., Increased levels of superoxide dismutase suppress meiotic segregation errors in aging oocytes. Chromosoma, 2019. 128(3): p. 215–222.

11. Anderson, L.K., et al., Juxtaposition of C(2)M and the transverse filament protein C(3)G within the central region of Drosophila synaptonemal complex. Proc Natl Acad Sci U S A, 2005. 102(12): p. 4482–7.

12. Buster, D.W., et al., SCFSlimb ubiquitin ligase suppresses condensin II-mediated nuclear reorganization by degrading Cap-H2. J Cell Biol, 2013. 201(1): p. 49–63.

13. Perkins, A.T. and S.E. Bickel, Using Fluorescence In Situ Hybridization (FISH) to Monitor the State of Arm Cohesion in Prometaphase and Metaphase I Drosophila Oocytes. J Vis Exp, 2017(130).

14. Perkins, A.T., et al., Oxidative stress in oocytes during midprophase induces premature loss of cohesion and chromosome segregation errors. Proc Natl Acad Sci U S A, 2016. 113(44): p. E6823–E6830.

15. Mahowald, A. and M. Kambysellis, Oogenesis, in The Genetics and Biology of Drosophila, M. Ashburner and T. Wright, Editors. 1980, Academic Press: New York. p. 141–224.

16. Spradling, A.C., Developmental Genetics of Oogenesis, in The development of Drosophila melanogaster, M. Bate and A. Martinez Arias, Editors. 1993, Cold Spring Harbor Laboratory Press: Cold Spring Harbor, NY. p. 1–70.

17. King, R.C., Ovarian development in Drosophila melanogaster. 1970, New York, NY: Academic Press. x, 227.

18. Khetani, R.S. and S.E. Bickel, Regulation of meiotic cohesion and chromosome core morphogenesis during pachytene in Drosophila oocytes. J Cell Sci, 2007. 120(Pt 17): p. 3123–37.

